# Cyclin Dependent Kinase 5 (CDK5) Regulates the Circadian Clock

**DOI:** 10.1101/728733

**Authors:** Andrea Brenna, Iwona Olejniczak, Rohit Chavan, Jürgen A. Ripperger, Sonja Langmesser, Elisabetta Cameroni, Zehan Hu, Claudio De Virgilio, Jörn Dengjel, Urs Albrecht

## Abstract

Circadian oscillations emerge from transcriptional and post-translational feedback loops. An important step in generating rhythmicity is the translocation of clock components into the nucleus, which is regulated in many cases by kinases. In mammals, the kinase promoting the nuclear import of the key clock component Period 2 (PER2) is unknown. Here we show that the cyclin-dependent kinase 5 (CDK5) regulates the mammalian circadian clock involving phosphorylation of PER2. Knock-down of *Cdk5* in the suprachiasmatic nuclei (SCN), the main coordinator site of the mammalian circadian system, shortened the free-running period in mice. CDK5 phosphorylated PER2 at serine residue 394 (S394) in a diurnal fashion. This phosphorylation facilitated interaction with Cryptochrome 1 (CRY1) and nuclear entry of the PER2-CRY1 complex. Taken together, we found that CDK5 drives nuclear entry of PER2, which is critical for establishing an adequate circadian period of the molecular circadian cycle. Therefore, CDK5 is critically involved in the regulation of the circadian clock and may represent a link to various diseases affected by the circadian clock.

## Introduction

The circadian clock, prevalent in most organisms, is an evolutionary adaptation to the daily light-dark cycle generated by the sun and the earth’s rotation around its own axis (Rosbash, 2009). This clock allows organisms to organize physiology and behavior over the 24 h time scale in order to adapt and thus optimize, body function to predictably recurring daily events. Malfunctioning or disruption of the circadian clock in humans results in various pathologies including obesity, cancer, and neurological disorders (Roenneberg & Merrow, 2016). In order to maintain phase synchronicity with the environmental light-dark cycle, the suprachiasmatic nuclei (SCN), a bipartite brain structure located in the ventral part of the hypothalamus above the optic chiasm, receive light information from the retina. The SCN convert this information into humoral and neuronal signals to set the phase of all circadian oscillators in the body (Dibner, Schibler, & Albrecht, 2010).

In order to measure the length of one day, organisms have developed cell-based molecular mechanisms relying on feedback loops involving a set of clock genes. The existence of such loops was suggested by the analysis of *Drosophila* having various mutations in their *period* (*per*) gene (Hardin, Hall, & Rosbash, 1990). Further studies completed the picture of intertwined transcriptional feedback loops at the heart of the *Drosophila* circadian oscillator (Darlington et al., 1998). Every day, per accumulates to a certain concentration upon which it enters into the nucleus together with timeless (tim). This protein complex inhibits transcriptional activation mediated by dClock and cycle acting on the expression of *per* and *tim*. After the degradation of the inhibitor complex, the repression is relieved and a new circadian cycle starts.

To fine-tune the period of the circadian oscillator, kinases regulate the accumulation and nuclear entry of per and tim. The kinase double-time (dbt) phosphorylates per to destabilize it and to prevent its transport into the nucleus (Kloss et al., 1998; Price et al., 1998). On the other hand, the kinase shaggy (shg) phosphorylates tim to stabilize the heterodimer and to promote its nuclear translocation (Martinek, Inonog, Manoukian, & Young, 2001). Many other kinases and phosphatases are necessary to complete the *Drosophila* circadian cycle and to adjust its phase to the external light-dark rhythm (Garbe et al., 2013).

The circadian oscillator of mammals is arranged very similarly to the one of *Drosophila*, with some modifications (Dibner et al., 2010; Takahashi, 2017). For instance, the function of *Drosophila* tim to escort per into the nucleus was replaced by the Cryptochromes (Cry) in the mammalian system (van der Horst et al., 1999). Furthermore, the first mutation to affect the mammalian circadian oscillator, *Tau*, was later mapped to Casein kinase Iε (CK1ε), which is the *Drosophila* dbt orthologue (Lowrey et al., 2000). One of the sites phosphorylated by CK1ε within human PER2 is mutated in the Familial Advanced Sleep Phase Syndrome (FASPS) (Toh et al., 2001). This mutation and also the *Tau* mutation were subsequently introduced into the mouse genome to prove their functional relevance (Meng et al., 2008; Y. Xu et al., 2007). However, a kinase similar to the function of shg in *Drosophila*, which stabilizes and promotes the import of PER proteins into the nucleus of mammals (Hirano, Braas, Fu, & Ptacek, 2017), has not been identified. Interestingly, PER2 contains over 20 potential phosphorylation sites (Vanselow et al., 2006), indicating that mammalian PER and specifically PER2 are highly regulated at the post-translational level. This degree of phosphorylation is probably contributing to the precise rhythmicity of PER2, which stands out as a crucial feature of the core clock (Chong, Ptacek, & Fu, 2012).

Among the plethora of kinases identified that phosphorylate mammalian clock proteins, cyclin dependent kinase 5 (CDK5) was found to target CLOCK (Kwak et al., 2013). CDK5 is a proline-directed serine-threonine kinase belonging to the Cdc2/Cdk1 family that is controlled by the neural specific activators p35, p39 (Tang et al., 1995; Tsai, Delalle, Caviness, Chae, & Harlow, 1994), and cyclin I (Brinkkoetter et al., 2009). CDK5 regulates various neuronal processes such as neurogenesis, neuronal migration, and axon guidance (Kawauchi, 2014). Outside of the nervous system CDK5 regulates vesicular transport, apoptosis, cell adhesion, and migration in many cell types (Contreras-Vallejos, Utreras, & Gonzalez-Billault, 2012). It has been proposed that CDK5 modulates the brain reward system (Benavides et al., 2007; Bibb et al., 2001) and that it is consequently linked to psychiatric diseases (Engmann et al., 2011; Zhu et al., 2012). Interestingly, the clock components PER2 and CLOCK have been associated with the same processes (Abarca, Albrecht, & Spanagel, 2002; Hampp et al., 2008; Roybal et al., 2007), leading us to speculate that an interaction between the circadian clock and CDK5 may exist. However, it is unknown whether CDK5 plays an important role in the central oscillator of the circadian clock.

In this study, we wanted to identify proteins promoting the nuclear transport of PER2 with focus on kinase(s) acting similarly to shg. Using a genetic synthetic lethal dosage screen in yeast, we observed a genetic interaction between *Per2* and *PHO85*, which encodes a cyclin-dependent protein kinase that is orthologous to CDK5 in mammals. Subsequent experiments in mice demonstrated that silencing of *Cdk5* in the SCN shortened the clock period. Our study identified CDK5 as a critical protein kinase in the regulation of the circadian clock and in particular as an important regulator of the crucial clock component PER2.

## Results

### Genetic interaction between Per2 and CDK5 in yeast and diurnal activity of CDK5

In order to gain insight into the regulation of PER2 function in mice, we initially tried to identify genes that genetically interact with *Per2* in budding yeast by using a variation of the Synthetic Genetic Array (SGA) method (Tong et al., 2001). To this end, we carried out a synthetic dosage lethality (SDL) screen, which is based on the concept that a high dosage of a given protein (*i.e.* PER2 in this case) may have negligible effect on growth in wild-type cells (as we found to be the case for PER2; Fig. 1A), but may compromise growth in mutants that have defects in pathway components or in functionally related processes (Measday et al., 2005; Sopko et al., 2006). Of note, SDL screens have been instrumental in the past to specifically predict the relationship between protein kinases and their targets (Sharifpoor et al., 2012). Our search in a yeast knockout collection (encompassing 4857 individual deletion strains) for mutants that exhibited significantly reduced growth when combined with increased dosage of PER2 (see Methods for further details) allowed us to isolate 3 mutants, namely *eap1*Δ, *gnd1*Δ, and *pho85*Δ (Fig. 1A). Among these, the strain lacking the cyclin-dependent protein kinase Pho85 was most dramatically compromised for growth in the presence of high doses of PER2. Hence, Pho85 antagonizes the growth-inhibitory effect of PER2 in yeast, which indicates that the Pho85-orthologous CDK5 may potentially act upstream of PER2 in mammalian cells.

**Figure 1:**
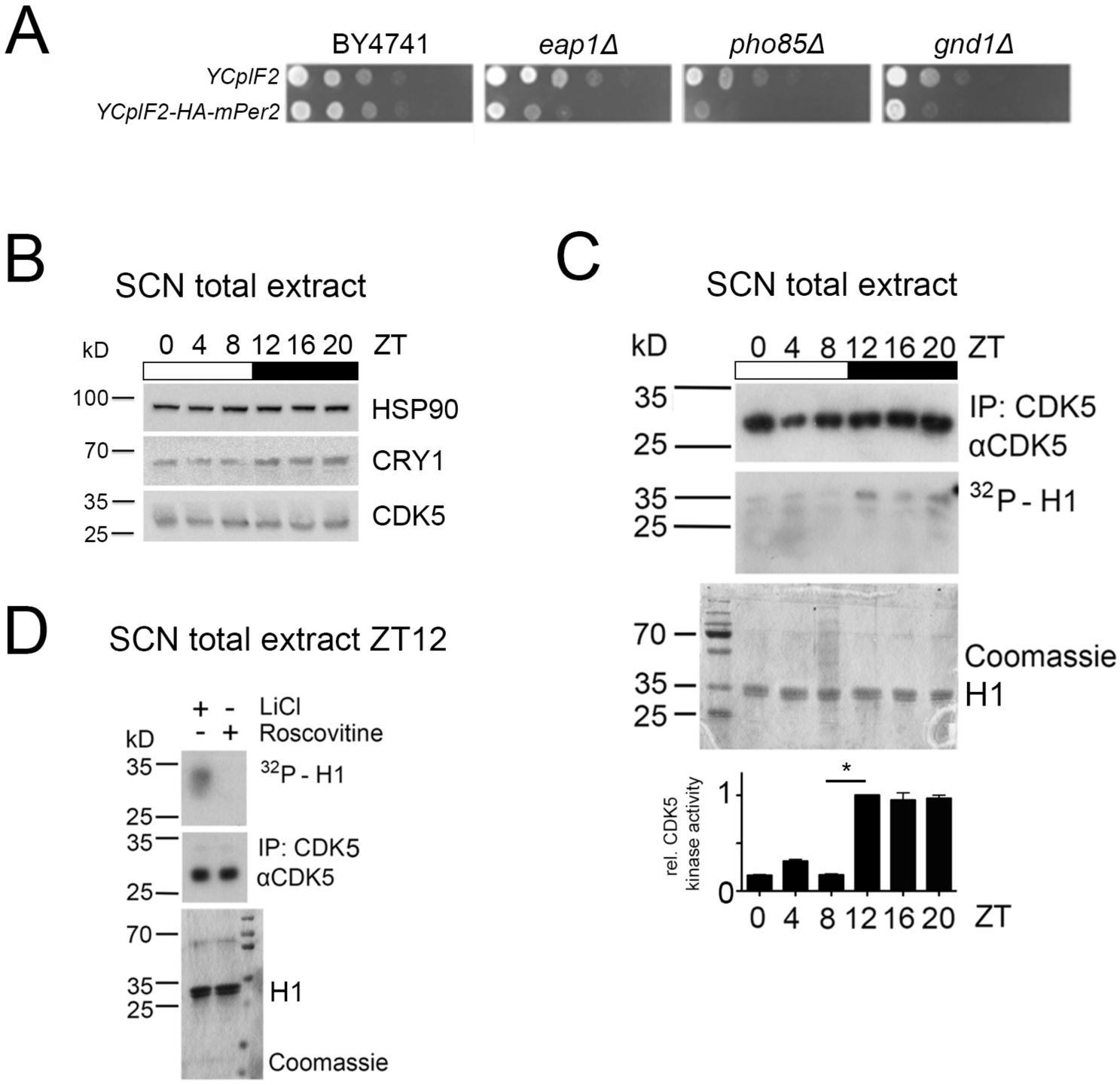
CDK5 intersects with PER2 and has diurnal activity in the SCN. (A) Loss of Eap1, Gnd1, or Pho85 compromises growth of PER2-overproducing yeast cells. The yeast mutants *eap1*Δ, *gnd1*Δ, and *pho85*Δ were identified in a synthetic dosage lethal screen as detailed under Methods. Wild-type (BY4741) as well as *eap1*Δ, *gnd1*Δ, and *pho85*Δ mutant cells carrying the control plasmid (YCpIF2) or the YPpIF2-*mPer2* plasmid (that drives expression of mouse PER2 from a galactose-inducible promoter) were pre-grown on glucose-containing SD-Leu media (to an OD_600_ of 2.0), spotted (in 10-fold serial dilutions) on raffinose and galactose-containing SD-Raf/Gal-Leu plates, and grown for 3 days at 30°C. (B) Immunoblot was performed on SCN extracts around the clock. SCN from seven animals were pooled at each indicated ZT between ZT0-20. Protein levels of CDK5, CRY1, and HSP90 were analyzed by Western Blot. (C) Diurnal activity of CDK5 was measured by an *in vitro* kinase assay. CDK5 was immunoprecipitated at each same time point between ZT0 and ZT20, and half of the immunoprecipitated material was used for performing an *in vitro* kinase assay using histone H1 (autoradiography, middle panel), whereas the other half was used to quantify the immunoprecipitated CDK5 (upper panel). Coomassie staining shows loading of the substrate (H1). Bottom panel: Quantification of 3 independent experiments (mean ± SEM). 1-way ANOVA with Bonferroni’s post-test, *: p<0.001. (D) The *in vitro* kinase assay was performed with SCN extracts at ZT12, and either LiCl (GSK3β inhibitor) or 34 µM roscovitine (CDK5 inhibitor). Histone H1 phosphorylation could not be detected with roscovitine treatment, showing the specificity of H1 phosphorylation by CDK5.

The protein kinase CDK5 is mostly expressed in the brain and has previously been implicated in phosphorylation of mammalian CLOCK (Kwak et al., 2013). However, the functional relevance of CDK5 for the clock mechanism has never been tested. Therefore, we investigated whether CDK5 affected the functioning of the circadian clock. First, we assessed whether CDK5 displayed time of day-dependent expression and activity in the SCN, the master clock of the circadian system. We collected SCN samples every 4 h starting from ZT0 until ZT20 (ZT0 = light on, ZT12 = light off), and performed western blots on total extracts using specific antibodies (Fig. 1B). The immunoblot against CRY1 showed a diurnal profile of this protein with a peak during the late-night phase, confirming that the mice were entrained properly to the light-dark cycle. In contrast, the CDK5 accumulation profile seemed to be unaffected by the time of day (Fig. 1B). Next, we investigated whether CDK5 kinase activity displayed a diurnal profile. While CDK5 levels did not change significantly over one day (Fig. 1B), we observed that histone-H1, a known CDK5 target (Peterson et al., 2010), was phosphorylated by this kinase in a time of day-dependent manner, with the highest levels of CDK5 activity observed at ZT12 to ZT20, i.e. during the dark phase (Fig. 1C). Phosphorylation of histone-H1 was specifically blocked by roscovitine, a CDK5 inhibitor (Hsu et al., 2013), whereas LiCl, a Gsk3β inhibitor, did not affect this phosphorylation (Fig. 1D), confirming a CDK5-specific phosphorylation. Altogether, these data demonstrated that CDK5 kinase activity (but not protein accumulation) was diurnal in the SCN.

### CDK5 regulates the circadian clock

Since CDK5 activity displayed a diurnal profile in the SCN, we tested whether knock-down of CDK5 in the master clock of the SCN changed circadian behavior in mice. To this end, we tested various shRNAs against *Cdk5* in NIH 3T3 fibroblast cells (Fig. S1A) and subsequently injected into the SCN region adeno-associated viral particles containing expression vectors for either a scrambled set of shRNA or a *Cdk5*-specific shRNA (variant D, Fig. S1A). After recovery from the procedure the animals were transferred into cages containing a running-wheel in order to assess their activity profiles. The control animals expressing the scrambled set of shRNA displayed normal activity in the light-dark (LD) cycle with precise onset of activity at the beginning of the dark phase (ZT12). This onset of activity was less precise in mice with a *Cdk5* knock-down (shCdk5) but comparable to animals with a deletion mutation in the clock gene *Per2*, designated as *Per2^Brdm1^* (Fig. 2A, Fig. S1B). In constant darkness (DD), χ^2^-periodogram analysis revealed a normal average free-running period for the scramble control mice, whereas for shCdk5 and *Per2^Brdm1^*, the period was significantly shortened (Fig. 2B). In one case, the shCdk5 animals became arrhythmic (Fig. 2C), again comparable to *Per2^Brdm1^* mice that eventually became arrhythmic in DD as well (Zheng et al., 1999). The total wheel-running activity was significantly reduced in shCdk5 and *Per2^Brdm1^* mice under DD as well as under LD conditions when compared with the scrambled control animals (Fig. S1C). The reduction of activity in the mutants under LD conditions is confined to the dark phase, but comparable between all three genotypes in the light phase (Fig. S1D). These results indicate that the period of the clock is affected by the lack of *Cdk5* expression in the SCN.

**Figure 2:**
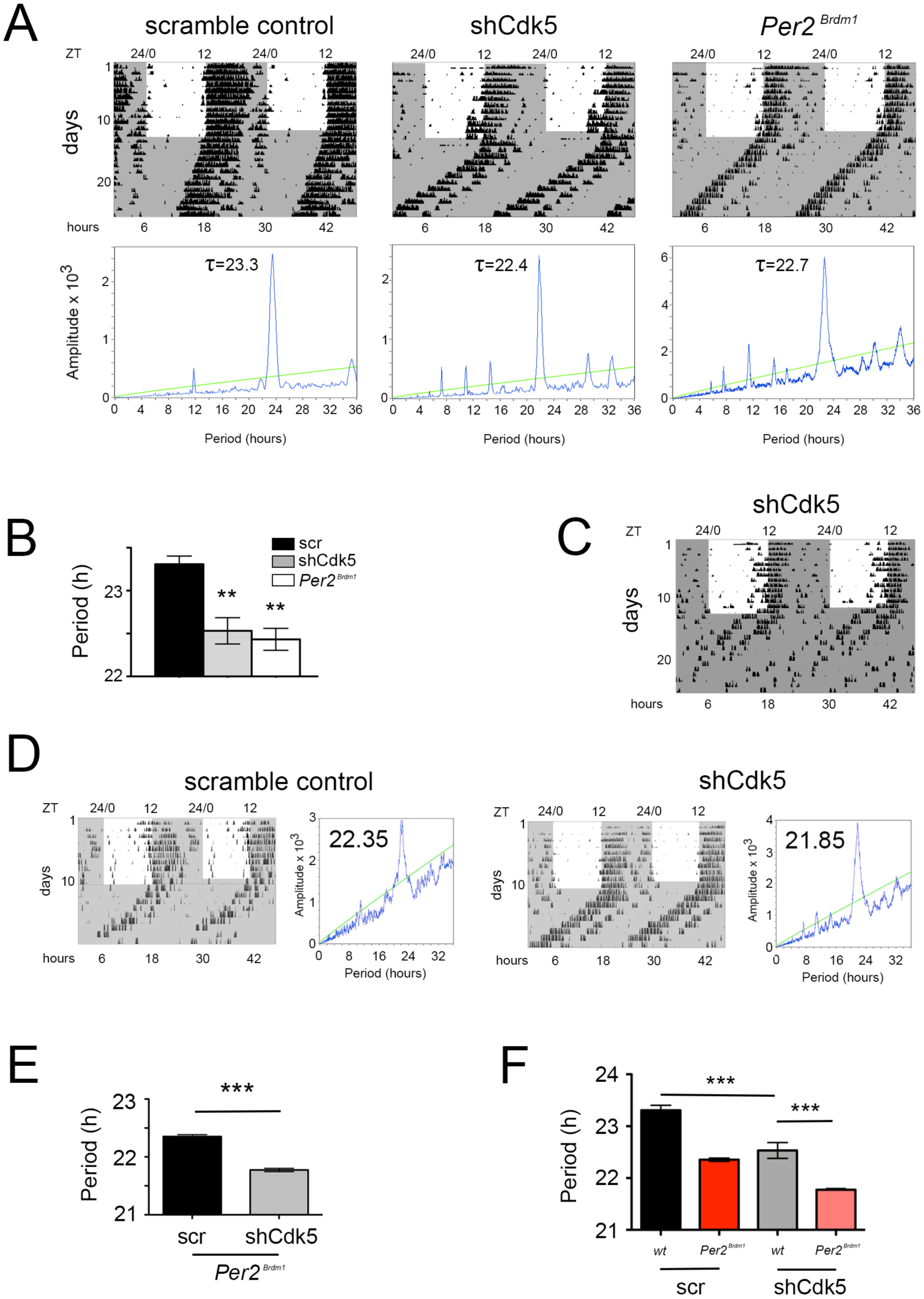
CDK5 affects the circadian clock. (A) Wheel-running activity of mice (black bins) infected with AAV expressing scrambled control shRNA, or shCdk5, and an animal with a deletion in the *Per2* gene (*Per2^Brdm1^*). The actograms are double plotted displaying in one row and below two consecutive days. The locomotor activity was confined to the dark period (shaded in grey), while under LD the mice displayed low activity during the light phase (white area). Under DD (continuous grey shaded area) the shCdk5 and *Per2^Brdm1^* animals show earlier onset of activity each day compared with the control animals. The χ^2^-periodogram analysis for each of the animals is shown below the corresponding actogram to determine the period length (τ). (B) Quantification of the circadian period: 23.3±0.1 h for the control mice (n=6, black bar), 22.5±0.2 h for shCdk5 injected mice (n=6, grey bar), and 22.4±0.1 h for *Per2^Brdm1^* mice (n=4, white bar), (mean±SEM). 1-way ANOVA with Bonferroni’s post-test, **p<0.01. (C) In some cases, mice in which *Cdk5* was silenced in the SCN became arrhythmic. (D) Wheel-running activity (black bins) of *Per2^Brdm1^* mice infected with AAV expressing scrambled control shRNA (scr), or shRNA against Cdk5 (shCdk5). The actograms are double plotted displaying in one row and below two consecutive days. The dark shaded area indicates darkness during which the free-running period was determined. To the right of each actogram the corresponding χ^2^-periodogram is shown. The number in each periodogram indicates the period of the animal. (E) Quantification of the circadian period: 22.35±0.03 h for the scrambled *Per2^Brdm1^* (n=3, black bar) and 21.77±0.03 h for the shCdk5 injected *Per2^Brdm1^* mice (n=5, grey bar). Values are the mean±SEM, t-test, ***p<0.0001. (F) 1-way ANOVA test on wild-type and *Per2^Brdm1^* animals infected with AAV expressing scrambled control shRNA (scr), or shRNA against Cdk5 (shCdk5). N = 3-6 animals, error bars are the mean±SEM, Bonferroni multiple comparisons test, ***p<0.001.

Interestingly, period in *Per2^Brdm1^* mutant and wild type shCdk5 knocked-down mice was not significantly different (Fig. 2F). In order to test the contribution of *Cdk5* we knocked down *Cdk5* in *Per2^Brdm1^* mutant mice. This even further shortened period in *Per2^Brdm1^* mutant animals compared to scramble control *Per2^Brdm1^* animals (Fig. 2D, E, S2), indicating that Cdk5 may affect period via other factors than *Per2*. Taken together, it appears that CDK5 is a main regulator of the circadian clock mechanism by targeting PER2.

In order to confirm that the different phenotypes were associated with the accumulation levels of CDK5 in control and *Cdk5*-silenced mice, we performed immunofluorescence assays on coronal sections of the SCN. Sections were stained with DAPI (blue) in order to label nuclei, with GFP antibody (green) in order to show cells infected by the virus, and with CDK5 antibody (red) in order to compare protein accumulation between the two strains. Scramble as well as shCdk5 mice expressed GFP in the SCN, indicating that the two different viruses infected cells in this brain region (Fig. 3A, S3A,B). The expression of *Cdk5* was efficiently suppressed in the SCN by the shCdk5 but not by the scrambled shRNA (Fig. 3A, S3A,B), indicating that the behavioral phenotypes observed are due to efficient knock-down of *Cdk5*. The *Cdk5* shRNAs was expressed in the SCN (the injection site) and to some extent also dorsal to the SCN but not in distant brain regions (i.e. the piriform cortex) as confirmed by lack of the GFP signal outside of the targeted region (Fig. 3A).

**Figure 3:**
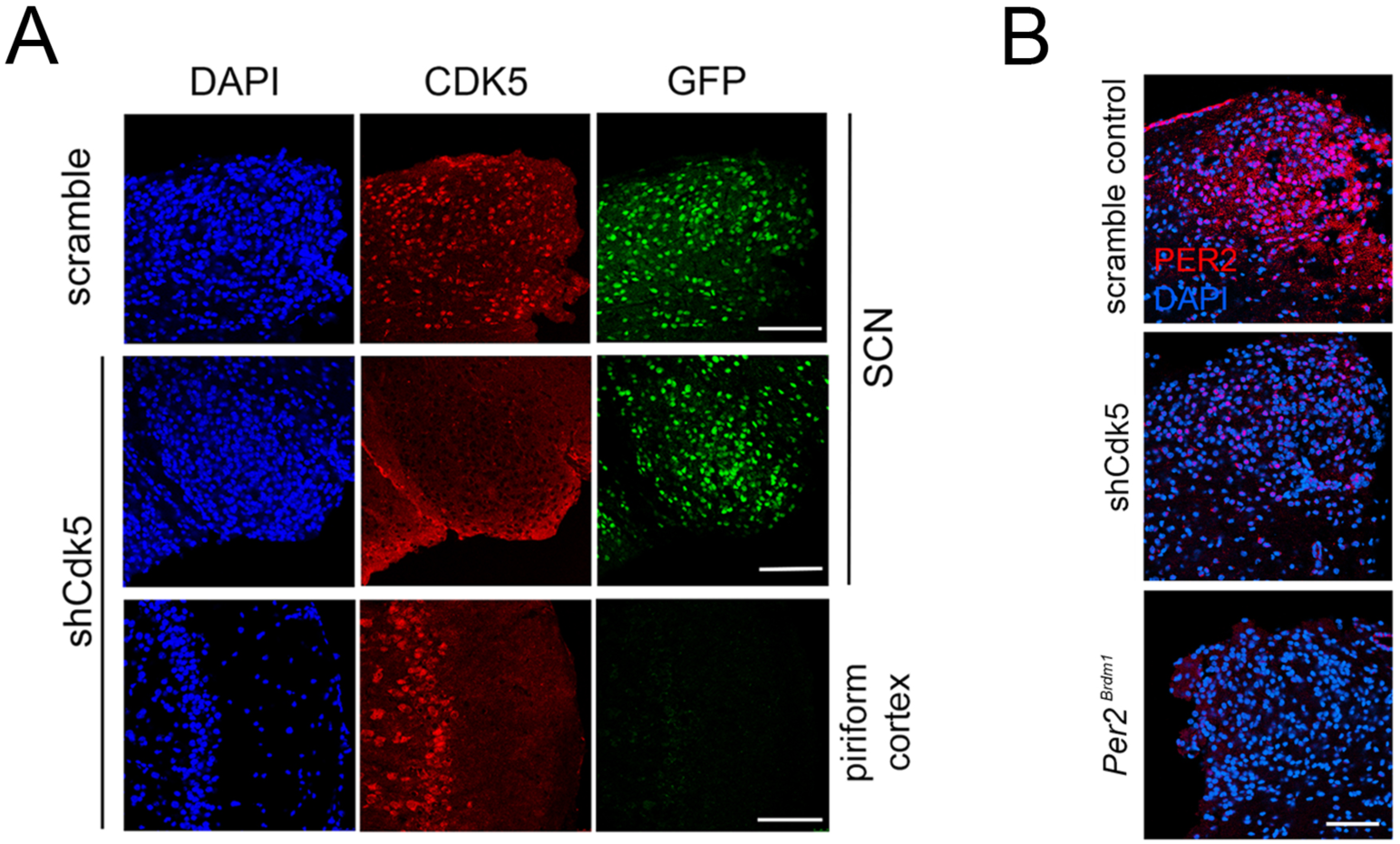
Immunohistochemistry in the SCN of control and shCdk5 silenced wild type and *Per2^Brdm1^* mice. (A) Representative sections of the SCN region after injection of AAVs carrying either scrambled shRNA, or shCdk5. Slices were stained with DAPI (blue), or anti-GFP (green) and anti-CDK5 (red) antibodies. GFP was used as marker for those cells infected by the virus. CDK5 was efficiently down-regulated in the SCN by shCdk5 (red panels) but not by scrambled shRNA, which was as efficiently delivered as shCDK5. As control, the non-infected piriform cortex from the same animal in which *Cdk5* was silenced is shown. Scale bar: 200 µm. (B) Analysis of PER2 expression in sections of the SCN of scrambled shRNA, shCdk5 and *Per2^Brdm1^* mice. Silencing of *Cdk5* leads to lack of PER2 (red) compared with control at ZT12, which almost resembles the situation observed in *Per2^Brdm1^* animals. Blue color: DAPI staining for cell nuclei. Scale bar: 200 µm.

Surprisingly, the phenotypes of shCdk5 and *Per2^Brdm1^* mice showed considerable similarity, implicating that the levels of PER2 accumulation might be similar in these two different mouse strains. In order to test whether *Cdk5* knock-down affected PER2, we stained with DAPI (blue) and immunostained with anti-PER2 (red) SCN sections obtained from control, shCdk5 and *Per2^Brdm1^* mice perfused at ZT12. PER2 was observed in the SCN of scramble controls, but was strongly reduced in shCdk5 and almost absent in *Per2^Brdm1^* animals (Fig. 3B, S3C,D). These data suggested that CDK5 is a main regulator of the core circadian clock in the SCN and may alter PER2 accumulation and potentially other proteins involved in clock regulation.

### CDK5 interacts with PER2 protein in a temporal fashion

A study in *Drosophila* has shown that several kinases, including cyclin dependent kinases, phosphorylate specific sites on per to maintain the circadian period (Garbe et al., 2013). Therefore, we aimed to understand whether a molecular interaction exists between CDK5 and PER2. We transfected cells with *Per2* and *Cdk5* expression vectors and tested whether the two proteins co-immunoprecipitated. We observed that immunoprecipitation with an anti-CDK5 antibody pulled down PER2 protein in two different cell lines (Fig. 4A, S4A). Similar interactions were observed when cells were transfected with expression constructs resulting in PER2 and CDK5 proteins fused to short amino-acid tags of viral protein 5 (V5) and haemaglutinine (HA) fused to them, respectively (Fig. 4B). Interestingly, interaction between PER2-V5 and CDK5-HA was reduced when roscovitine, which inhibits interaction of CDK5 with its targets (Hsu et al., 2013), was added to the cells (Fig. 4B). This suggested that active CDK5 protein interacted better with PER2 than CDK5 in its inhibited form.

**Figure 4:**
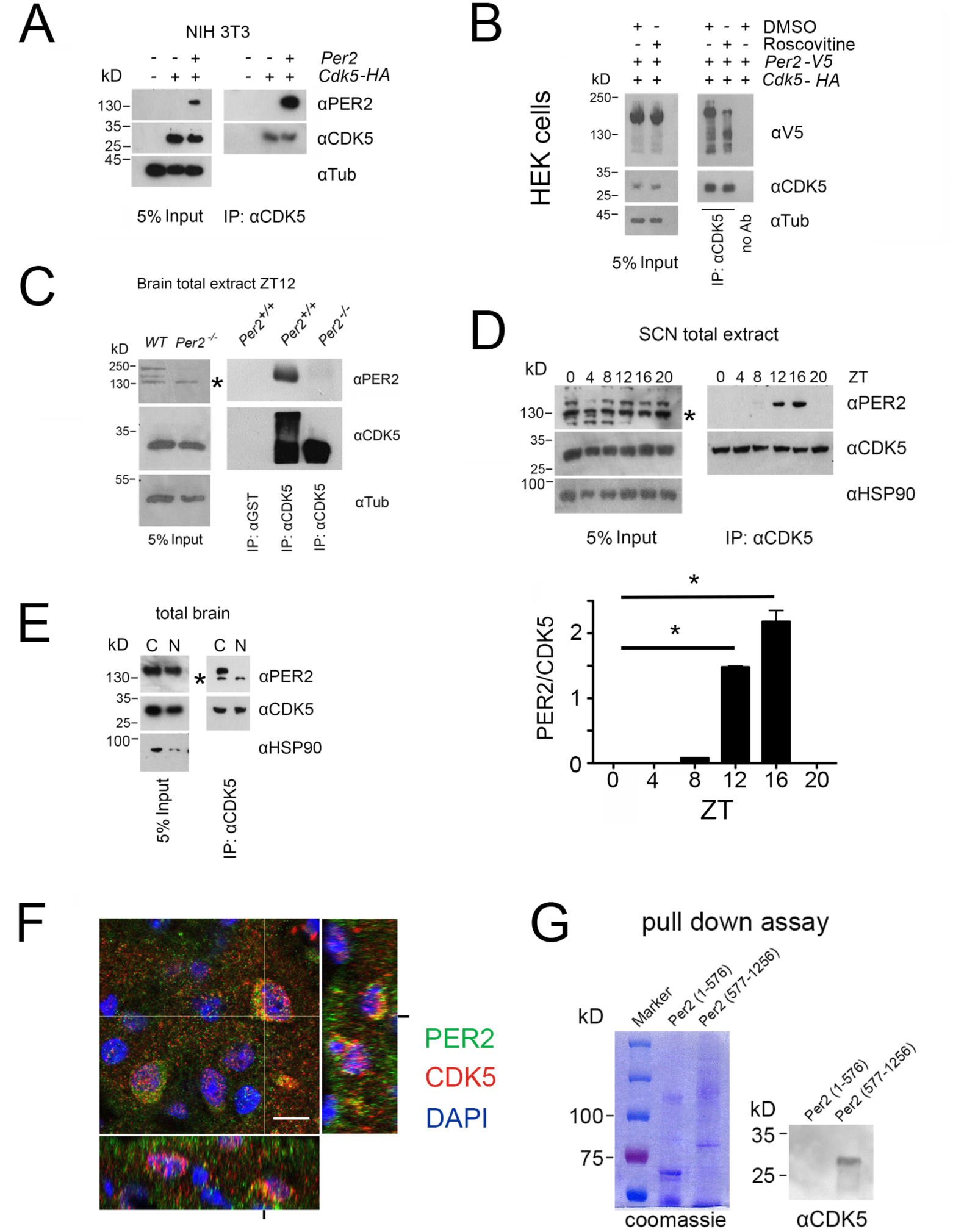
PER2 interacts with CDK5 in a temporal fashion in the cytoplasm. (A) Overexpression of PER2 and CDK5-HA in NIH 3T3 cells and subsequent immunoprecipitation (IP) using an anti-CDK5 antibody. The left panel shows 5% of the input and the right panel co-precipitation of PER2 with CDK5. (B) Overexpression of PER2-V5 and CDK5-HA in HEK293 cells in presence or absence of 34 µM roscovitine (CDK5 inhibitor) and DMSO (solvent). Left panel shows 5% of the input and the right panel the immunoprecipitation with anti-CDK5 or without antibody. (C) Immunoprecipitation (IP) of PER2 and CDK5 from total mouse brain extract collected at ZT12. Left panel shows the input. The right panel depicts co-immunoprecipitation of PER2 and CDK5 using either anti-CDK5 antibody or anti-GST antibody for precipitation. The middle lane shows PER2-CDK5 co-immunoprecipitation in control animals (*Per2^+/+^*) but not in *Per2^−/−^* mice illustrating the specificity of the PER2-CDK5 interaction. The * in the blot indicates unspecific signal. (D) Temporal profile of the PER2-CDK5 interaction in total extracts from SCN tissue around the clock. Input was analyzed by immunoblot using anti-CDK5, anti-PER2, and anti-HSP90 antibodies (left panel). CDK5 co-immunoprecipitated PER2 in a diurnal fashion with a peak between ZT12 and ZT16. The statistical analysis of the PER2/CDK5 signal around the clock is shown below (1-way ANOVA with Bonferroni’s post-test, n=3, *p<0.0001, values are mean ± SEM). * in the blot indicates unspecific signal. (E) Immunoprecipitation of PER2 with CDK5 from cytoplasmic and nuclear brain extracts collected at ZT12. The left panel shows the input and the right panel co-IP of PER2 and CDK5, which occurs only in the cytoplasm but not in the nucleus. The smaller band detected by the anti-PER2 antibody depicts an unspecific band that is smaller than PER2. * in the blot indicates unspecific signal. (F) Slices from the SCN obtained at ZT12 were immunostained with PER2 antibody (green), CDK5 (red), and nuclei were marked with DAPI (blue). Co-localization of the two proteins results in the yellow color. Scale bar: 10 µm. The z-stacks right and below the micrograph confirm co-localization of PER2 and CDK5 (yellow). (G) Purification of the N-terminal half of PER2 (1-576) or the C-terminal half (577-1256) (left panel, coomassie). CDK5-His was pulled down by both recombinant PER2 attached to the glutathione resin, but only the C-terminal was able to retain CDK5 (immunoblot using anti-His antibody, right panel).

In order to test whether this interaction could be observed in tissue, we prepared total brain extracts at ZT12, when kinase activity of CDK5 was high (Fig. 1C). At two different salt concentrations we could pull-down PER2 and CDK5 using either anti-CDK5 or anti-PER2 antibodies (Fig. S4B). The specificity of the signals was confirmed by using brain extracts from *Per2^−/−^* mice (Chavan et al., 2016) that completely lack PER2 protein (Fig. 4C). Next, we wanted to investigate whether the interaction between the two proteins is time of day-dependent in the SCN. Total extracts of SCN tissue at ZT0, 4, 8, 12, 16 and 20 were prepared and immunoprecipitation with an anti-CDK5 antibody pulled down PER2 at ZT8, 12, and 16, with the strongest signals at ZT12 and ZT16 (Fig. 4D). Taken together, these observations suggested that the interaction between CDK5 and PER2 can occur in brain tissue and that in the SCN this interaction was time of day-dependent. This observation was confirmed on SCN tissue sections, where we observed PER2 expression at ZT12 but less at ZT0 with co-localization of CDK5 restricted to ZT12 (Fig. S4C).

Next, we tested in which subcellular compartment the interaction between CDK5 and PER2 takes place. We prepared nuclear and cytoplasmic extracts from total brain tissue and performed immunoprecipitation using an anti-CDK5 antibody. PER2 could only be observed in the cytoplasmic but not the nuclear fraction (Fig. 4E). This was supported by the observation that the two proteins were co-localized only in the cytoplasm in SCN tissue (Fig. 4F, yellow color).

Furthermore, we evaluated with which part of PER2 the CDK5 protein interacts. We tested whether deletions in the PAS-domain of PER2, a known domain for protein interactions (Ponting & Aravind, 1997), influenced CDK5 binding. No significant effect of deletions of the PAS-A and PAS-B domains on the interaction was observed (Fig. S4D). Next, we generated expression vectors coding either for the N-terminal (1-576) or the C-terminal part (577-1257) of PER2 fused to GST (Fig. S4E). The recombinant forms of PER2 and histidine-tagged CDK5 were produced in bacteria. A pull-down assay with these proteins showed that the C-terminal but not the N-terminal half of the PER2 protein was pulled-down by CDK5, suggesting that CDK5 binds to the C-terminal part of PER2 (Fig. 4G). This does, however, not exclude weak interactions of the CDK5 protein with the N-terminal half *in vivo*. Taken together, our data suggest a physical interaction of PER2 and CDK5 in the cytoplasm.

### CDK5 phosphorylates PER2 at serine 394

In order to understand whether CDK5 phosphorylates the PER2 protein we overexpressed the N-terminal and C-terminal parts of PER2 fused to GST in bacteria (Fig. S5A) and performed an *in vitro* kinase assay with the recombinant proteins. Recombinant CDK5/p35 protein complex along with γ-^32^P labeled ATP resulted in phosphorylation of the N-terminal part of the PER2 protein with a main signal at around 120 kD (Fig. 5A, S5B, ^32^P panels). Addition of roscovitine abolished phosphorylation of PER2 whereas LiCl had no effect (Fig. S5C). Interestingly, no phosphorylation of the C-terminal part of PER2 was observed, only a signal corresponding to the auto-phosphorylation of CDK5 was detected at around 60 kD (Fig. 5A, ^32^P panel).

**Figure 5:**
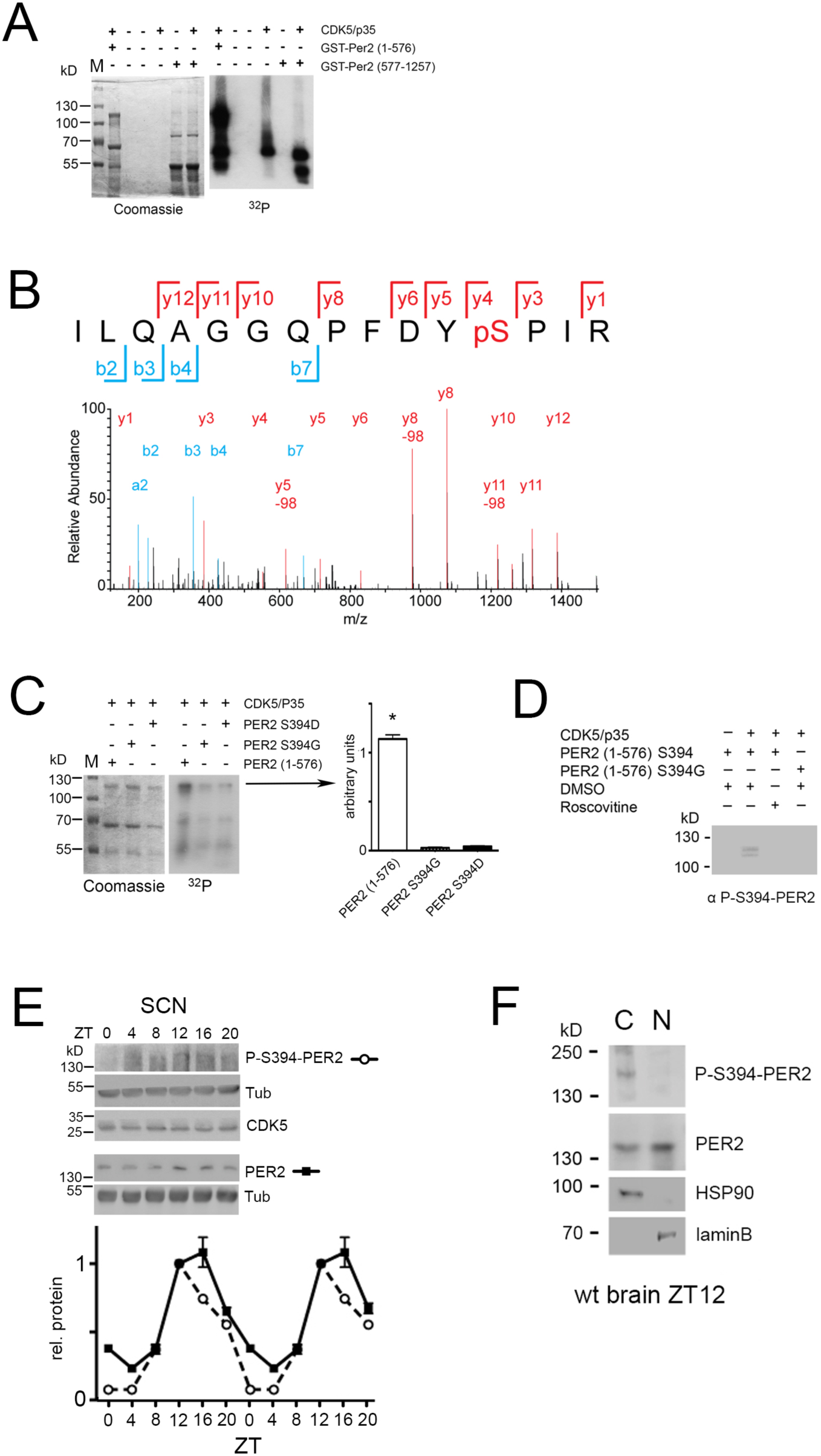
CDK5 phosphorylates PER2 at S394. (A) An *in vitro* kinase assay was performed using recombinant CDK5/p35 and either GST-PER2 1-576 or GST-PER2 577-1256 as substrate. The samples were subjected to 10% SDS page (Coomassie, left panel) and the phosphorylation of PER2 was detected by autoradiography in order to visualize ^32^P-labeled proteins (right panel). CDK5 phosphorylates the N-terminal half (1-576) of a GST-PER2 fusion protein whereas the C-terminal half (577-1257) is not phosphorylated. The signal for CDK5/p35 alone indicates CDK5 auto-phosphorylation seen in all lanes when CDK5 is present. (B) Annotated mass spectrum of the tryptic peptide PER2^383-397^ ILQAGGQPFDYpSPIR containing the phosphorylated residue S394. The red color depicts the y-ion series (1-12) and blue the b-ion series (2-7, a2); y5-98, y8-98, y11-98 show the de-phosphorylated ions. (C) *In vitro* kinase assay was performed as in (A). The putative phosphorylation site was mutated to aspartic acid (S394D) or glycine (S394G). Both mutations abrogated the CDK5-mediated phosphorylation. Coomassie staining reveals equal expression of the GST-PER2 fragments. The bar diagram at the right shows the quantification of 3 experiments. 1-way-ANOVA with Bonferroni’s post-test, *: p<0.001 (D) The monoclonal antibody produced against P-S394-PER2 does recognizes PER2 (1-576) S394 phosphorylation mediated by CDK5/p35 in presence but not in absence of the kinase or when CDK5 is inactivated by roscovitine. This antibody does not recognize the S394G mutated form even in presence of CDK5/p35. (E) Temporal profile of P-S394-PER2 and total PER2 in SCN tissue. Upper panels show Western blots of the corresponding proteins indicated on the right. Below the quantification of 3 experiments is shown, in which the value at ZT12 of PER2 has been set to 1. The data were double plotted. Values are the mean±SEM. 2-way ANOVA with Bonferroni’s multiple comparisons revealed that the two curves are significantly different with p < 0.0001, F=93.65, DFn=1, DFd=48. (F) Subcellular localization of P-S394-PER2. Total wild-type mouse brain extracts were separated into cytoplasmic (HSP90 positive) and nuclear (laminB positive) fractions. Phosphorylated PER2 was predominantly detected in the cytoplasm with the P-S394-PER2 antibody, whereas the general PER2 antibody detected PER2 in both compartments with higher amounts in the nuclear fraction.

Next, we aimed to identify the phosphorylation site(s) in the N-terminal part of PER2 using the recombinant protein, which was phosphorylated by CDK5/p35 *in vitro*. Mass spectrometry revealed several phosphorylation sites at serine and threonine residues, respectively (Supplemental Table S1). One of the serine residues of PER2 was located within a CDK5 consensus sequence and had the highest probability score for being phosphorylated (Fig. 5B). The serine residue at position 394 (S394) of PER2 is located at the end of the PAS domain and within the deletion of the mutated PER2 of *Per2^Brdm1^* mice (Zheng et al., 1999). This suggested that CDK5/p35 phosphorylates S394 and that this phosphorylation is of functional relevance. Mutations of this serine to aspartic acid (S394D) or glycine (S394G) reduced phosphorylation by CDK5/p35 significantly (Fig. 5C), confirming that CDK5/p35 phosphorylated S394. Next, we produced a monoclonal antibody against the phosphorylated serine at 394 of PER2 (P-S394-PER2) (Fig. S5D-F). With this antibody we detected the phosphorylated N-terminal fragment of PER2 in presence of CDK5/p35 but not when S394 was mutated to glycine (S394G) or when CDK5 was inhibited by roscovitine (Fig. 5D), confirming S394 phosphorylation by CDK5/p35.

In order to determine whether PER2 phosphorylation at S394 is time of day-dependent, we collected SCN tissue every 4 h. The P-S394-PER2 specific antibody detected highest phosphorylation at ZT12 with weaker or no phosphorylation at other time points indicating that S394 is phosphorylated in a time of day-dependent manner (Fig. 5E). Fractionation of wild-type brain cellular extracts prepared at ZT12 into nuclear and cytoplasmic parts showed phosphorylated S394 predominantly in the cytoplasm with little or no signal in the nucleus when labeled with the P-S394-PER2 antibody (Fig. 5F). Total PER2 was observed in both cellular compartments with higher levels in the nucleus (Fig. 5F). This suggested that phosphorylation of S394 of PER2 happens predominantly in the cytoplasm and that this phosphorylation is either removed or occluded when PER2 enters the nucleus.

### CDK5 affects stability and nuclear localization of PER2

To evaluate the function of CDK5-driven PER2 phosphorylation we wanted to determine whether CDK5 affects PER2 stability. We treated NIH 3T3 cells with roscovitine and DMSO as control and determined endogenous levels of PER2. We observed that roscovitine treatmetn of cells reduced PER2 levels, suggesting that CDK5 can affect protein stability (Fig. 6A). In order to challenge this observation, we deleted *Cdk5* in NIH 3T3 cells using the CRISPR/Cas9 method (Fig. S6A-C). We observed that deletion of *Cdk5* led to reduced amounts of PER2 (Fig. 6B), consistent with the data shown in Figure 6A. These observations support the notion that phosphorylation by CDK5 affects PER2 abundance. In order to monitor PER2 stability, we knocked down *Cdk5* using the shRNA D (Fig. S1A). We observed that increasing amounts of shCdk5 dampened PER2 levels proportionally to the decreasing CDK5 levels (Fig. 6C).

**Figure 6:**
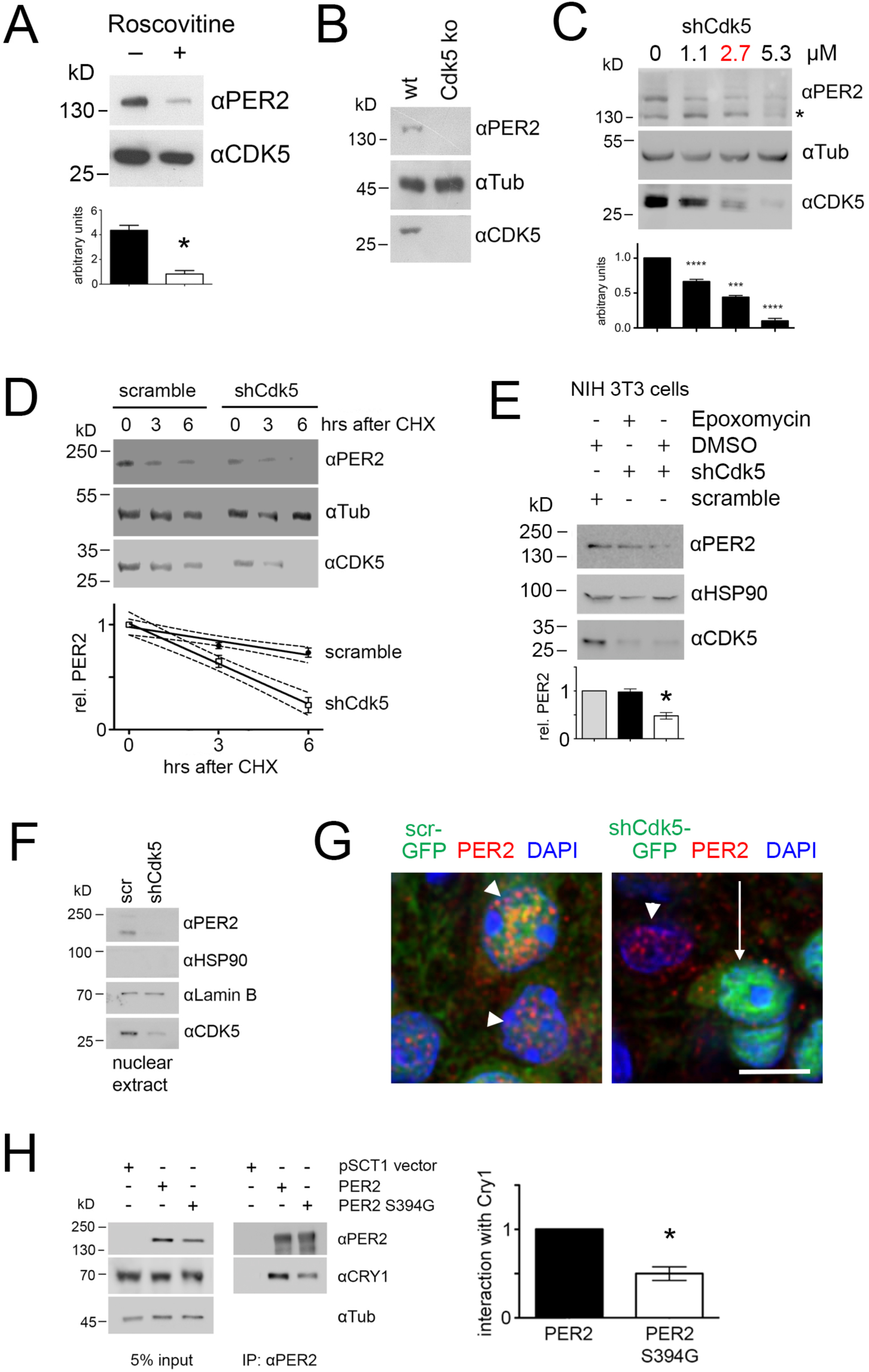
CDK5 affects PER2 stability and nuclear localization. (A) Western blot of NIH 3T3 cell extracts with and without roscovitine treatment. When roscovitine inhibited CDK5, less PER2 protein was detected in cell extracts. The bar diagram below shows values (mean±SEM) of 3 experiments with significant differences between roscovitine treated and untreated cells, t-test, *p < 0.001. (B) CRISPR/Cas9-mediated knockout of *Cdk5* in NIH 3T3 cells. Western blot shows absence of PER2 in cells when *Cdk5* is deleted. (C) Titration of CDK5 knock-down as revealed by Western blotting. PER2 levels decreased proportionally to increasing amounts of shCdk5. 2.7 µM of shCdk5 (red) was used for subsequent experiments. The value without shCdk5 was set to 1. 1-way ANOVA with Bonferroni post-test, n=4, ***p<0.001, ****p<0.0001, mean±SD. The * in the blot indicates unspecific signal. (D) Temporal profile of protein abundance in NIH 3T3 cells 0, 3 and 6 h after inhibition of protein synthesis by 100 µM cycloheximide (CHX) in presence of scrambled shRNA, or shCdk5, respectively (2.7 µM of the respective shRNA was used). The diagram below shows quantification of PER2 protein over time. Linear regression with 95% confidence intervals (hatched lines) indicates that knock-down of *Cdk5* leads to less stable PER2 (shCdk5 t_1/2_=4h, scr t_1/2_=11h). 2-way ANOVA with Bonferroni’s post-test revealed that the two curves are significantly different, n=3, p<0.01, F=24.53, DFn=1, DFd=4. (E) Inhibition of the proteasome by epoxomycin in cells with shCdk5 leads to amounts of PER2 that are higher compared with the levels without epoxomycin treatment and are comparable to the levels observed in cells without *Cdk5* knockdown. Diagram below displays the quantification of 3 experiments. Scrambled shRNA values were set to 1. 1-way ANOVA with Bonferroni’s post-test shows no significant reduction of PER2 in shCdk5 cells in presence of epoxomycin, but significantly lower values in absence of epoxomycin when compared with scrambled shRNA treatment. 1-way ANOVA with Bonferroni’s post-test, n=3, p<0.001. (F) PER2 abundance in nuclear extracts of NIH 3T3 cells. Knockdown of *Cdk5* reduces PER2 levels in the nucleus as revealed by Western blotting. HSP90 = cytosolic marker, LaminB = nuclear maker. (G) Immunofluorescence of PER2 (red) at ZT12 in mouse SCN sections after infection with AAV (green) expressing scrambled shRNA (left panel), or shCdk5 (right panel). Nuclei are visualized by DAPI staining (blue). PER2 can only be observed in the nucleus in presence (white arrow heads) but not in absence of CDK5 (white arrow). Scale bar = 7.5 µm. (H) Co-immunoprecipitation of CRY1 by PER2 in NIH 3T3 cells. Substitution of S394 to G in PER2 reduces the levels of co-precipitated CRY1 (right panel). The left panel shows the input. The bar diagram on the right displays the quantification of 3 experiments, where the amount of precipitated CRY1 by PER2 is set to 1. Paired t-test reveals a significant difference between the amounts of CRY1 precipitated by PER2 and the S394G PER2 mutation, n = 3, *p<0.05, mean±SD.

In order to determine whether CDK5 modulates degradation of PER2, we blocked protein synthesis using cycloheximide. Under conditions that partially knocked down *Cdk5* (at a concentration of 2.7 µM of shCdk5, Fig. 6C), we measured PER2 and CDK5 protein levels over 6 h after cycloheximide treatment. We found that degradation of PER2 was faster when *Cdk5* was knocked down compared with unspecific shRNA treatment (shCdk5 t_1/2_=4 h, scr t_1/2_=11 h) (Fig. 6D), indicating that reduction of *Cdk5* accelerated PER2 degradation. Next, we investigated whether PER2 degradation involved the proteasome. Cells were treated with epoxomycin, a proteasome inhibitor, or with the solvent DMSO. In line with our previous experiments, shCdk5 treatment efficiently knocked down CDK5 and reduced PER2 levels compared with scrambled shRNA treatment. Addition of epoxomycin, but not DMSO, significantly increased PER2 levels despite absence of CDK5 (Fig. 6E), indicating that PER2 degradation involved the proteasome. Residual amounts of CDK5 in the cells still may phosphorylate PER2 and direct it into the nucleus. Therefore, we wanted to see whether PER2 could be detected in nuclear extracts of shCdk5 knocked down cells. In line with our previous observations we did not detect PER2 in nuclear extract (Fig. 6F), supporting the idea that PER2 needed to be phosphorylated by CDK5 in order to enter the nucleus. Data from immunofluorescence experiments on SCN sections were in line with this hypothesis. PER2 was only detected in nuclei when CDK5 was available (Fig. 6G, arrowheads, S6D), but not when shCdk5 was expressed in SCN cells (Fig. 6G, white arrow, S6D).

It has been described that nuclear entry of PER2 involves CRY1 (Kume et al., 1999; Ollinger et al., 2014). In addition, CRY1-mediated hetero-dimerization stabilizes PER2 by inhibiting its own ubiquitination (Yagita et al., 2000). Therefore, we tested the interaction potential of wild-type PER2 and the S394G PER2 mutation with CRY1 by overexpressing the two PER variants in NIH 3T3 cells. Immunoprecipitation of wild-type PER2 pulled down CRY1; however, the S394G PER2 mutation was significantly less efficient in doing so (Fig. 6H). The small amounts of CRY1 detected may be bound to endogenous PER2 that is present in the cells. In summary, these experiments suggested that CDK5 affects PER2 stability, interaction with CRY1, and nuclear localization.

## Discussion

Not only do kinases play a crucial role in signal transduction in response to extracellular stimuli, but they also regulate cycling processes such as the cell cycle and circadian rhythms. Most cyclin dependent kinases (CDKs) regulate the cell cycle, with few exceptions such as the cyclin dependent kinase 5 (CDK5). This kinase is ubiquitously expressed and its function is vital in post-mitotic neurons, where other CDKs are not active. Although CDK5 is not implicated in cell cycle progression, it can aberrantly activate components of the cell cycle when it is dysregulated in post-mitotic neurons, leading to cell death (Chang, Vincent, & Shah, 2012). Interestingly, cell death is affected by the clock component PER2 as well (Magnone et al., 2014), suggesting that both, CDK5 and PER2 act in the same pathway, or that their pathways cross at a critical point during the regulation of cell death. The synthetic dosage lethal screen that we performed in yeast supports this notion, as expression of PER2 in yeast lacking *Cdk5* strongly and significantly compromised growth (Fig. 1A).

The kinase CDK5 displays many effects that ensure proper brain function and development. Mice deficient for *Cdk5* are perinatal lethal (Gilmore, Ohshima, Goffinet, Kulkarni, & Herrup, 1998; Ohshima et al., 1996). CDK5 influences cortical neuron migration, cerebellar development, synapse formation and plasticity (Kawauchi, 2014). Here, we identified a new role for this kinase, i.e. the regulation of the circadian clock *in vivo*. Previously, CDK5 had been identified to phosphorylate CLOCK and thereby regulate CLOCK stability and cellular distribution in cells (Kwak et al., 2013). In the SCN, however, NPAS2 may replace the function of CLOCK (Debruyne et al., 2006; DeBruyne, Weaver, & Reppert, 2007) and therefore phosphorylation of CLOCK by CDK5 may play a minor role in the SCN. Hence, to unravel the novel function of CDK5 in the circadian oscillator, we had to restrict ourselves to the use of SCN tissue and whole animals.

CDK5 activity, but not its protein accumulation, displays a diurnal profile in the SCN with high activity during the night and low activity during the day (Fig. 1C). The activity displayed a typical on/off profile similar to other CDKs. This finding raises the question how this diurnal activity of CDK5 may be achieved. On one hand, ATP accumulation, which is required for phosphorylation, peaks during the night in the SCN (Yamazaki, Maruyama, Cagampang, & Inouye, 1994). On the other hand, CDK5 activity is regulated by cofactors. Depending on its cofactor, CDK5 in the brain phosphorylates targets involved in neurodegenerative diseases (e.g. Tau, MAP1B), neuronal migration (e.g. DCX), and synaptic signaling (e.g. Ca_v_2.2, Dynamin1, NR2A, DARPP-32) (Kawauchi, 2014). The most obvious candidates to regulate its time-dependent activity would be cyclins D1 and E, which inhibit CDK5, or cyclin I, which activates it. Alternatively, other known CDK5 regulators such as p35 may be involved (Shah & Lahiri, 2014). Most likely, positive and negative feedback loops of other kinases and phosphatases are necessary to generate the on/off profile, although the components involved in this mechanism are probably different from the ones known for CDKs that regulate the cell cycle. Interestingly, CK1 phosphorylates and activates CDK5 *in vitro* (Sharma, Sharma, Amin, Albers, & Pant, 1999) and CDK5 phosphorylates and inhibits CK1δ (Ianes et al., 2016) establishing a feedback loop between the two kinases. However, additional research is needed to determine the precise mechanism of diurnal on/off activation of CDK5.

CDK5 binds to the C-terminal half of PER2 (Fig. 4G) and phosphorylates it at S394 (Fig. 5), which is located in the PAC domain of the N-terminal half of the protein. Hence, the binding and phosphorylation sites are far apart, suggesting a structure of PER2 allowing proximity of the CDK5 binding and phosphorylation domains. We cannot exclude weak binding of CDK5 to the N-terminal half of PER2, because phosphorylation at S394 occurs *in vitro* even in the absence of the C-terminal half of the PER2 protein (Fig. 5A). This may be due to the fact that the N-terminal half is overexpressed *in vitro*, which strongly increases the probability of phosphorylation by CDK5 even in the absence of the C-terminal binding domain. It is also known that p35 (which is used in the *in vitro* kinase assay to activate CDK5) can increase the interaction between CDK5 and its targets (Hsu et al., 2013).

In SCN tissue PER2 phosphorylation at S394 appears to be time of day-dependent, with highest levels at ZT12 and ZT16 (Fig. 5E) when CDK5 activity is high (Fig. 1C). Compared with total PER2 protein the S394 phosphorylated form appears to be slightly advanced in its phase. The difference in phase is probably even larger than it appears here, because the polyclonal antibody that detects total PER2 also detects the phosphorylated S394 PER2 variant. This is especially important in the rise of the signal detected, which appears to be identical in figure 5E. Probably the steep increase between ZT8 and ZT12 represents the S394 phosphorylated forms in both curves. In contrast, the decrease in PER2 levels differs between total PER2 and P-S394-PER2 form. Consistent with previous studies total PER2 peaks in the nucleus at ZT16 in the SCN (Nam et al., 2014) when P-S394-PER2 is not detected anymore. This highlights that additional post-translational modifications of PER2 exist (Toh et al., 2001; Vanselow et al., 2006) and that P-S394-PER2 disappears faster compared with other modified forms. Probably, P-S394-PER2 plays a role in PER2 dynamics in terms of shuttling from the cytoplasm to the nucleus, because P-S394-PER2 can only be observed in the cytoplasmic and not the nuclear fraction (Fig. 5F). The phosphorylation of PER2 by CDK5 may therefore be critical for the assembly of a macromolecular complex in the cytoplasm (Aryal et al., 2017), which then enters the nucleus.

The difference in the decline between PER2 and its S394 phosphorylated form in the SCN may suggest a role of the S394 phosphorylation not only for nuclear transport but also for PER2 protein stability. The earlier decline of the P-S394-PER2 signal compared with total PER2 (Fig. 5F) might suggest that the S394 phosphorylated form is less stable. Apparently, the opposite is the case, as shown in Fig. 6. Pharmacological inhibition of CDK5 (Fig. 6A), CRISPR/Cas9 mediated knock-out of *Cdk5* (Fig. 6B), and shRNA mediated knock-down of *Cdk5* (Fig. 6C) all led to reduced levels of PER2 in cells. The half-life of PER2 is clearly increased in the presence of CDK5, rising from about 4 h to 11 h, indicating that phosphorylation at S394 has a stabilizing function. This is in accordance with previous results that described almost absent levels of PER2 in the *Per2^Brdm1^* mutant mice (Zheng et al., 1999). This mouse strain expresses a PER2 lacking 87 amino acids in the PAS and PAC domains, where the S394 and the CDK5 consensus sequence are localized. CDK5 cannot phosphorylate this mutant PER2 and therefore the protein is less stable. As a consequence, the formation of the macromolecular complex responsible for nuclear transport of PER2 is disturbed. This results in a temporal change of BMAL1/CLOCK/NPAS2 activity, shortening the clock period. Accordingly, *Per2^Brdm1^* mutant mice display a short period or no circadian period in constant darkness (Zheng et al., 1999), similar to the phenotype observed for the CDK5 knock-down mice (Fig. 2B).

PER2 stability is affected by CK1δ/ε, which phosphorylate PER2 at several sites and regulate degradation of PER2 via the proteasome (Eide et al., 2005; Y. Xu et al., 2007; Narasimamurthy et al., 2018). This effect is similar to the action of dbt on *Drosophila* per. Interestingly, CDK5 can phosphorylate CK1δ to reduce its activity (Ianes et al., 2016). This phosphorylation could cross-regulate the activities of both kinds of kinases to fine-tune the amount of PER2. This is evidenced by the observation, that knock-down of *Cdk5* in *Per2^Brdm1^* mutant mice further shortens period in these animals (Fig. 2D, E). The mammalian orthologue of shg, Gsk3β, does not phosphorylate the mammalian Tim but the nuclear receptor NR1D1 (Mukherji, Kobiita, & Chambon, 2015). This change in substrate may be related to the shift in function of the CRYs to replace Tim in the mammalian circadian oscillator. Similar to shg, CDK5 phosphorylation of PER2 increases its half-life (Fig. 6D). Lack of CDK5, and therefore lack of phosphorylation at S394 of PER2, leads to proteasomal degradation of PER2 as evidenced by epoxomycin treatment, which inhibits the proteasome and reduces the decline of PER2 levels in the cell (Fig. 6E). This is consistent with a recent report that describes the ubiquitin ligase MDM2 as controlling PER2 degradation via the proteasome (Liu et al., 2018). However, it is not clear whether it is the phosphorylation at S394 *per se* or the capacity to participate in a macromolecular complex to enter the nucleus that stabilizes PER2. In any case, this phosphorylation appears to be essential for nuclear entry of PER2 (Fig. 6F, G).

A recent report showed that mammalian PER represses and de-represses transcription by displacing BMAL1-CLOCK from promoters in a CRY-dependent manner (Chiou et al., 2016). Our data support these findings. PER2 containing a S394G mutation, which abolishes CDK5-mediated phosphorylation, displayed reduced interaction potential with CRY1 (Fig. 6H). Because CRY1 is involved in nuclear transport of PER2 (Kume et al., 1999; Ollinger et al., 2014; Yagita et al., 2000), lack of interaction with the S394G mutant form of PER2 leaves this protein in the cytoplasm, unable to enter the nucleus (Fig. 6G). The present data are also in agreement with previous experiments in which we investigated the role of protein phosphatase 1 (PP1) and its effects on the circadian clock (Schmutz et al., 2011). Expression of a specific PP1 inhibitor in the brain lengthened circadian period and increased PER2 levels and its nuclear accumulation in neurons. These effects are all opposite to what we observe when PER2 is not phosphorylated at S394. Therefore, it could be speculated that PP1 is involved in the dephosphorylation of P-S394, thereby counterbalancing phosphorylation of this site by CDK5.

Taken together, our results indicate that CDK5 phosphorylates PER2 at S394. This phosphorylation appears to be important for PER2 to bind efficiently to CRY1 in order to allow entry of PER2 into the nucleus. Inhibition of CDK5 in cells leads to degradation of PER2 in the proteasome (Fig. 7). Inhibition of CDK5 *in vivo* inhibits nuclear entry of PER2 and shortens period to a similar extent as observed in *Per2^Brdm1^* mutant mice, which express a barely detectable level of protein lacking 87 amino acids including S394. Taken together, CDK5 regulates the circadian clock and influences PER2 nuclear transport via phosphorylation. Because PER2 is involved in several physiologically relevant pathways in addition to clock regulation (Albrecht, Bordon, Schmutz, & Ripperger, 2007), PER2 may mediate several biological functions that were previously linked to CDK5, such as the regulation of the brain reward system (Benavides et al., 2007; Bibb et al., 2001) and psychiatric diseases (Engmann et al., 2011; Zhu et al., 2012).

**Figure 7.**
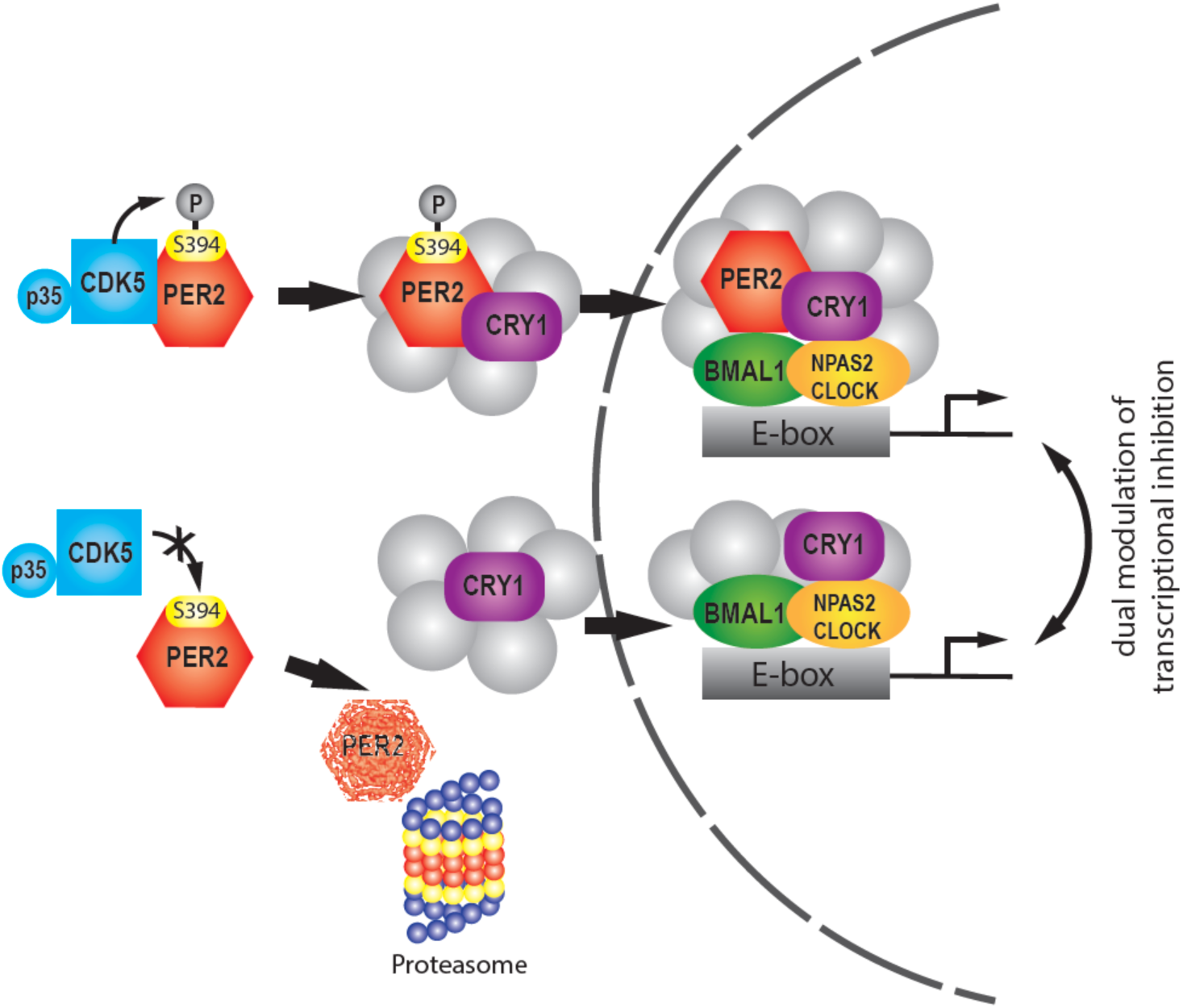
Model showing the regulation of PER2 by CDK5. The upper row illustrates phosphorylation of PER2 at S394 by CDK5 that subsequently favors interaction with CRY1 and leads to transport into the nucleus, where the PER2/CRY1 complex inhibits BMAL1/NPAS2 (or in the periphery CLOCK)-driven transcriptional activation. The lower part illustrates that inhibition of CDK5 leads to a lack of S394 PER2 phosphorylation, which renders the PER2 protein more prone to degradation by the proteasome. CRY1 does not form a complex with PER2 and hence PER2 is not transported into the nucleus. CRY1 enters the nucleus independently and can inhibit the BMAL1:NPAS2 (or in the periphery CLOCK) transcriptional complex. This model is consistent with the dual modulation of transcriptional inhibition (Ye et al., 2014; H. Xu et al., 2015). Transcriptional inhibition is modulated in an intricate unknown manner by various additional factors (grey) (Aryal et al., 2017) that may be cell type specific.

## Methods

### Animals and housing

All mice were housed with food and water *ad libidum* in transparent plastic cages (267 mm long × 207 mm wide × 140 mm high; Techniplast Makrolon type 2 1264C001) with a stainless-steel wire lid (Techniplast 1264C116), kept in light- and soundproof ventilated chambers. All mice were entrained to a 12:12-h LD cycle, and the time of day was expressed as Zeitgeber time (ZT; ZT0 lights on, ZT12 lights off). Two- to four-month-old males were used for the experiments. Housing as well as experimental procedures were performed in accordance with the guidelines of the Schweizer Tierschutzgesetz and the declaration of Helsinki. The state veterinarian of the Canton of Fribourg approved the protocol. The floxed Per2 mice (Chavan et al., 2016) are available at the European Mouse Mutant Archive (EMMA) strain ID EM:10599, B6;129P2-Per2^tm1Ual^/Biat.

### Synthetic dosage lethal (SDL) screen

The SDL screen was essentially performed as described earlier (Measday et al., 2005; Tong et al., 2001). Briefly, the bait strain Y2454 (MATα *mfa1Δ::MFA1pr-HIS3, can1Δ, his3Δ1, leu2Δ0, ura3Δ0, lys2Δ0*) carrying the plasmid YCplF2-*mPer2* (that drives expression of PER2 from the galactose-inducible *GAL1* promoter) was inoculated into 50 mL glucose-containing synthetic dropout medium lacking leucine (SD-Leu) and grown at 30°C overnight with shaking. Cells were then centrifuged, resuspended in 20 mL of the supernatant, poured into a sterile rectangular petri dish, spotted in a 96-well format on rectangular SD-Leu plates (coined “bait plates” hereafter) using a Biomek 2000 robot (Beckman Coulter, USA), and then grown at 30° C for three days. In parallel, the gene deletion array in the strain BY4741 (MATa *his3*Δ*1*, *leu2*Δ*0*, *met15*Δ*0*, *ura3*Δ*0*) was spotted from the storage plates onto fresh G418-containing YPD plates (96-well format) and also grown at 30°C for three days. For the mating procedure (overnight at 30°C), colonies from bait plates were (robotically) spotted onto plates containing YPD (plus adenine) and the colonies from the gene deletion array plates were (each separately and in duplicate) spotted on top of them. The next day, the colonies were transferred to G418-containing SD plates lacking lysine, methionine, and leucine (SD-Lys/Met/Leu/+G418) to select for diploids that harbour the YCplF2-*mPer2* plasmid. Diploids were then spotted onto plates containing sporulation medium (10 g L^−1^ potassium acetate, 1g L^−1^ yeast extract, 0.1 g L^−1^ glucose, 2% w/v agar, supplemented with uracil, histidine, and G418) and incubated at 24°C. After 9 days, tetrads were observed and the colonies were transferred to canavanine-containing SD plates lacking arginine and histidine (SD-Arg/His/+canavanine) to select for MATa haploids. Following growth at 30°C for three days, a second haploid selection was carried out by spotting the colonies on SD-Arg/His/Leu/+canavanine plates (to select for MATa haploids containing the YCplF2-*mPer2* plasmid). Following growth at 30°C for two days, a third haploid selection was carried out by spotting cells on SD-Arg/His/Leu/+canavanine/+G418 plates (to select for MATa haploids containing the YCplF2-*mPer2* plasmid as well as the respective gene deletions of the yeast knockout collection). Following incubation at 30°C for five days, colonies were then spotted in parallel onto SD-Arg/His/Leu/+G418 plates and on SD-Raf/Gal-Arg/His/Leu/+G418 plates (containing 1% raffinose and 2% galactose as carbon sources) to induce expression of PER2. Both types of plates were incubated at 30°C for four days and photographed every day. Strains that grew significantly less well on SD-Raf/Gal-Arg/His/Leu/+G418 than on SD-Arg/His/Leu/+G418 included *eap1*Δ, *gnd1*Δ, and *pho85*Δ. In control experiments, the respective original yeast knockout collection mutants were transformed in parallel with the YCplF2-*mPer2* or the empty YCplF2 plasmid (Foreman & Davis, 1994), selected on SD-Leu plates, grown overnight in liquid SD-Leu, spotted (10-fold serial dilutions) on SD-Raf/Gal-Leu plates, and grown for 3 days at 30° (Figure 1A). Please note that all media containing G418 were made with glutamate (1 g L^−1^) instead of ammonium sulfate as nitrogen source, as recommended in (Tong et al., 2001).

### Adeno Associate Virus (AAV) production and stereotaxic injections

Adeno Associate Viruses (AAVs) were produced in the Viral Vector Facility (ETH Zurich). Plasmids used for the production are available on the VVF web site. Two constructs were produced. ssAAV-9/2-hSyn1-chI[mouse(shCdk5)]-EGFP-WPRE-SV40p(A) carried the shRNA against Cdk5 (shD, see Fig. S1A or Supplemental Table 2) which knocked down only neuronal *Cdk5*. ssAAV-9/2-hSyn1-chI[1x(shNS)]-EGFP-WPRE-SV40p(A) was the scrambled control.

Stereotaxic injections were performed on 8-week-old mice under isofluorene anaesthesia using a stereotaxic apparatus (Stoelting). The brain was exposed by craniotomy and the Bregma was used as reference point for all coordinates. AAVs were injected bilaterally into the SCN (Bregma: anterior-posterior (AP) − 0.40 mm; medial-lateral (ML) ± 0.00 mm; dorsal-ventral (DV) – 5.5 mm, angle +/− 3°) using a hydraulic manipulator (Narishige: MO-10 one-axis oil hydraulic micromanipulator, http://products.narishige-group.com/group1/MO-10/electro/english.html) at a rate of 40 nL/min through a pulled glass pipette (Drummond, 10 µl glass micropipet; Cat number: 5-000-1001-X10). The pipette was first raised 0.1mm to allow spread of the AAVs, and later withdrawn 5 min after the end of the injection. After surgery, mice were allowed to recover for 2 weeks and entrained to LD 12:12 prior to behavior and molecular investigations.

### Locomotor activity monitoring

Analysis of locomotor activity parameters was done by monitoring wheel-running activity, as described in (Jud, Schmutz, Hampp, Oster, & Albrecht, 2005), and calculated using the ClockLab software (Actimetrics). Briefly, for the analysis of free-running rhythms, animals were entrained to LD 12:12 and subsequently released into constant darkness (DD). Internal period length (τ) was determined from a regression line drawn through the activity onsets of ten days of stable rhythmicity under constant conditions. Total and daytime activity, as well as activity distribution profiles, was calculated using the respective inbuilt functions of the ClockLab software (Acquisition Version 3.208, Analysis version 6.0.36). Numbers of animals used in the behavioral studies are indicated in the corresponding figure legends.

### Immunofluorescence

Animals used for the immunohistochemistry were killed at appropriate ZTs indicated in the corresponding figure legends. Brains were perfused with 0.9% NaCl and 4% PFA. Perfused brains were cryoprotected by 30% sucrose solution and sectioned (40 µm, coronal) using a cryostat. Sections chosen for staining were placed in 24-well plates (2 sections per well), washed three times with 1x TBS (0.1 M Tris/0.15 M NaCl) and 2x SSC (0.3 M NaCl/0.03 M tri-Na-citrate pH 7). Antigen retrieval was performed with 50% formamide/2x SSC by heating to 65°C for 50 min. Then, sections were washed twice in 2x SSC and three times in 1x TBS pH 7.5, before blocking them for 1.5 h in 10% fetal bovine serum (Gibco)/0.1% Triton X-100/1x TBS at RT. After the blocking, the primary antibodies, rabbit anti-PER2-1 1:200 (Alpha Diagnostic, Lot numb. 869900A1.2-L), mouse anti-Cdk5 clone 2H6 1:20 (Origene, Lot numb. A001), and rabbit anti-GFP 1:500 (abcam ab6556) diluted in 1% FBS/0.1% Triton X-100/1x TBS, were added to the sections and incubated overnight at 4°C. The next day, sections were washed with 1x TBS and incubated with the appropriate fluorescent secondary antibodies diluted 1:500 in 1% FBS/0.1% Triton X-100/1x TBS for 3 h at RT. (Alexa Fluor 488-AffiniPure Donkey Anti-Rabbit IgG (H+L) no. 711–545–152, Lot: 132876, Alexa Fluor647-AffiniPure Donkey Anti-Mouse IgG (H+L) no. 715–605–150, Lot: 131725, Alexa Fluor647-AffiniPure Donkey Anti-Rabbit IgG (H+L) no. 711–602–152, Lot: 136317 and all from Jackson Immuno Research). Tissue sections were stained with DAPI (1:5000 in PBS; Roche) for 15 min. Finally, the tissue sections were washed again twice in 1x TBS and mounted on glass microscope slides. Fluorescent images were taken by using a confocal microscope (Leica TCS SP5), and images were taken with a magnification of 40x or 63x. Images were processed with the Leica Application Suite Advanced Fluorescence 2.7.3.9723 according to the study by Schnell et al. (Schnell et al., 2014).

Immunostained sections were quantified using ImageJ version 1.49. Background was subtracted and the detected signal was divided by the area of measurement. An average value obtained from three independent areas for every section was used. The signal coming from sections obtained from silenced mice was quantified as relative amount to the scramble, which was set to 1. Statistical analysis was performed on 3 animals per treatment.

### Cell culture

NIH3T3 mouse fibroblast cells (ATCCRCRL-1658™) were maintained in Dulbecco’s modified Eagle’s medium (DMEM), containing 10% fetal calf serum (FCS) and 100 U/mL penicillin-streptomycin at 37°C in a humidified atmosphere containing 5% CO2. Cdk5 KO cells were produced applying the CRISPR/Cas9 technique according to the manufacturer’s protocol of the company (Origene, SKU # KN303042).

### Plasmids

The following plasmids used were previously described: pSCT-1, pSCT-1mPer2, pSCT-1 mPer-V5, pSCT1 ΔPasA mPer2 -V5, pSCT1 ΔPasB mPer2 -V5 (Langmesser, Tallone, Bordon, Rusconi, & Albrecht, 2008) (Schmutz, Ripperger, Baeriswyl-Aebischer, & Albrecht, 2010). pSCT-1 Cdk5-HA, pet-15b Cdk5-HIS, Gex-4T Per2 1-576, pGex-4T Per2 577-1256 were produced for this paper. The full-length cDNA (or partial fragments) encoding PER2 and the full-length Cdk5 were previously sub-cloned in the TOPO vector according to the manufacturer’s protocol (Catalog numbers pCR™2.1-TOPO® vector: K4500-01). They were subsequently transferred into the plasmid pSCT-1 using appropriate restriction sites. pGex-4T Per2 1-576 S394G, S394D, pSCT-1 mPer2 S394G were obtained using site-specific mutagenesis according to the manufacturer’s protocol on the requested codon carrying the interested amino acid of interest (Agilent Catalog # 200518). For accession numbers, vectors, mutations, and primers sources, see Supplemental Table 2.

### Transfection and co-immunoprecipitation of overexpressed proteins

NIH 3T3 cells were transfected in 10 cm dishes at about 70% of their total confluency using linear polyethylenimine (LINPEI25; Polysciences Europe). The amounts of expression vectors were adjusted to yield comparable levels of expressed protein. Thirty hours after transfection, protein extracts were prepared. With regard to immunoprecipitation, each antibody mentioned in the paper was used in the conditions indicated by the respective manufacturer. The next day, samples were captured with 50 µL at 50% (w/v) of protein-A agarose beads (Roche) at 50% (w/v) and the reaction was kept at 4° C for 3 h on a rotary shaker. Prior to use, beads were washed 3 times with the appropriate protein buffer and resuspended in the same buffer (50% w/v). The beads were collected by centrifugation and washed 3 times with NP-40 buffer (100 mM Tris-HCl pH7.5, 150 mM NaCl, 2 mM EDTA, 0.1% NP-40). After the final wash, beads were resuspendend in 2% SDS, 10% glycerol, 63 mM Trish-HCL pH 6.8 and proteins were eluted for 15 min at RT. Laemmli buffer was finally added, samples were boiled for 5 min at 95° C and finally loaded onto 10% SDS-PAGE gels (Laemmli, 1970).

### Total protein extraction from cells (Ripa method)

Medium was aspirated from cell plates, which were washed 2 times with 1x PBS (137mM NaCl, 7.97 mM Na_2_HPO_4_ x 12 H_2_O, 2.68 mM KCl, 1.47 mM KH_2_PO_4_). Then PBS was added again and plates were kept 5 min at 37°C. NHI3T3 or HEK cells were detached and collected in tubes and washed 2 times with 1x PBS. After the last washing, pellets were frozen in liquid N_2_, resuspended in Ripa buffer (50 mM Tris-HCl pH7.4, 1% NP-40, 0.5% Na-deoxycholate, 0.1 % SDS, 150 mM NaCl, 2 mM EDTA, 50 mM NaF) with freshly added protease or phosphatase inhibitors, and homogenized by using a pellet pestle. After that samples were centrifuged for 15 min at 16’100 g at 4° C. The supernatant was collected in new tubes and pellet discarded.

### Total protein extraction from brain tissue

Total brain or isolated SCN tissue was frozen in liquid N_2,_ and resuspended in lysis buffer (50 mM Tris-HCl pH 7.4, 150 mM NaCl, 0.25% SDS, 0.25% sodium deoxycholate, 1 mM EDTA) and homogenized by using a pellet pestle. Subsequently, samples were kept on ice for 30 min and vortexed 5 times for 30 sec each time. The samples were centrifuged for 20 min at 12’000 rpm at 4° C. The supernatant was collected in new tubes and the pellet discarded.

### Nuclear/cytoplasmic fractionation

Tissues or cells were resuspended in 100 mM Tris-HCl pH 8.8/10 mM DTT and homogenized with a disposable pellet pestle. After 10 min incubation on ice, the samples were centrifuged at 2’500 g for 2 min at 4°C and the supernatant discarded. After adding 90 µL of completed cytoplasmic lysis buffer (10 mM EDTA, 1 mM EGTA, 10 mM Hepes pH 6.8, 0.2% Triton X-100, protease inhibitor cocktail (Roche), NaF, PMSF, ß-glycerophosphate), the pellet was resuspended by vortexing, followed by centrifugation at 5’200 rpm for 2 min at 4°C. The supernatant transferred into a fresh 1.5 mL tube was the CYTOPLASMIC EXTRACT. The pellet was washed three times with cytoplasmic lysis buffer and resuspended in 45 µL 1x NDB (20% glycerol, 20 mM Hepes pH 7.6, 0.2 mM EDTA, 2 mM DTT) containing 2x proteinase and phosphatase inhibitors followed by adding 1 volume of 2x NUN (2 M Urea, 600 mM NaCl, 2% NP-40, 50 mM Hepes pH 7.6). After vortexing the samples were incubated 30 min on ice, centrifuged 30 min at 13’000 rpm at 4°C and the supernatant that was transferred into a fresh tube was the NUCLEAR EXTRACT.

### Immunoprecipitation using brain tissue extracts

A protein amount corresponding to between 400 and 800 µg of total extract was diluted with the appropriate protein lysis buffer in a final volume of 250 µL and immunoprecipitated using the indicated antibody (ratio 1:50) and the reaction was kept at 4° C overnight on a rotary shaker. The day after, samples were captured with 50 µL of 50% (w/v) protein-A agarose beads (Roche) and the reaction was kept at 4° C for 3 h on a rotary shaker. Prior to use, beads were washed 3 times with the appropriate protein buffer and resuspended in the same buffer (50% w/v). The beads were collected by centrifugation and washed 3 times with NP-40 buffer (100 mM Tris-HCl pH7.5, 150 mM NaCl, 2 mM EDTA, 0.1% NP-40). After the final wash, beads were resuspendend in 2% SDS 10%, glycerol, 63 mM Trish-HCL pH 6.8 and proteins were eluted for 15 min at RT. Laemmli buffer was finally added, samples were boiled 5 min at 95° C and loaded onto 10% SDS-PAGE gels.

### Pull-down assay with GST-Per2 fragments

GST-fused recombinant Per2 proteins were expressed in the *E. coli* Rosetta strain [plasmids: GST-Per2 N-M (1-576), GST-Per2 M-C (577-1256)]. Proteins were induced with 1 mM IPTG (Sigma-Aldrich) for 3 h at 30°C. Subsequently, proteins were extracted in an appropriate GST lysis buffer (50 mM Tris-Cl pH 7.5, 150 mM NaCl, 5% glycerol) adjusted to 0.1% Triton X-100 and purified to homogeneity with glutathione-agarose beads for 2 h at 4°C. The beads were then incubated overnight at 4° C and washed with GST lysis buffer adjusted with 1 mM DTT. Subsequently, elution with 10 mM reduced glutathione took place for 15 min at room temperature. Elution was stopped by adding Laemmli buffer and samples were loaded onto the gel after 5 min at 95° C and WB was performed using anti-GST (Sigma no. 06-332) and anti-HA antibodies (Roche no. 11867423001) for immunoblotting.

### CRISPR/Cas9 *Cdk5* knock-out cell line

The CRISPR/Cas9 Cdk5 cell line was produced starting from NIH3T3 cells using a kit provided by Origene (www.origene.com). The knock-out cell line was produced according to the manufacturer’s protocol. Briefly, cells at 80% of confluency were co-transfected with a donor vector containing the homologous arms and functional cassette, and the guide vector containing the sequence that targets the *Cdk5* gene. In parallel, a scrambled negative guide was also co-transfected with a donor vector. 48 h after transfection the cells were split 1:10 and grown for 3 days. Cells were split another 7 times (this time is necessary to eliminate the episomal form of donor vector, in order to have only integrated forms). Then, single colonies were produced and clones were analyzed by PCR in order to find those clones that did not express *Cdk5* RNA. Positive clones were re-amplified.

PCR primers for genomic Cdk5:
FW: 5’-tgtgagtaccacctcctctgcaa-3’
RW: 5’-ttaaacaggccaggcccc-3’

### Knockdown of Cdk5

About 24 h after seeding cells, different shRNA Cdk5 plasmids (Origene TL515615 A/B/C/D Cdk5 shRNA) were transfected to knock down *Cdk5* according to the manufacturer’s instructions. The knock-down efficiency was assessed at 48 h post transduction by Western blotting. Scrambled shRNA plasmid (Origene TR30021) was used as a negative control.

### Cycloheximide treatment

NIH3T3 cells were treated with 100 µM cycloheximide 48 h after transfection with the indicated vectors, and cells were harvested 0, 3, and 6 h after treatment.

### Proteasome inhibitor treatment

About 48 h after transfection with either scrambled or shCdk5, cells where *Cdk5* was silenced were treated for 12 h with either DMSO (vehicle) or epoxomicin (Sigma-Aldrich) at a final concentration of 0.2 µM. Samples were collected, and proteins extracted followed by Western blotting.

### *In vitro* kinase assay

Recombinant GST-fused PER2 protein fragments were expressed and purified from the BL21 Rosetta strain of *E. coli* according to the manufacturer’s protocol described before, using glutathione-sepharose affinity chromatography (GE Healthcare). Each purified protein (1 µg) was incubated in the presence or absence of recombinant Cdk5/p35 (the purified recombinant N-terminal His6-tagged human Cdk5 and N-terminal GST-tagged human p25 were purchased from Millipore). Reactions were carried out in a reaction buffer (30 mM Hepes, pH 7.2, 10 mM MgCl2, and 1 mM DTT) containing [γ-^32^P] ATP (10 Ci) at room temperature for 1 h and then terminated by adding SDS sample buffer and boiling for 10 min. Samples were subjected to SDS-PAGE, stained by Coomassie Brilliant Blue, and dried, and then phosphorylated proteins were detected by autoradiography.

### *In vitro* kinase assay using immunoprecipitated Cdk5 from SCN

CDK5 was immunoprecipitated from SCN samples at different ZTs (circa 600 µg of protein extract) (Fig. S7). After immunoprecipitation at 4° C overnight with 2x Protein A agarose (Sigma-Aldrich), samples were diluted in washing buffer and split in two halves. One half of the IP was used for an in vitro kinase assay. Briefly, 1 µg of histone H1 (Sigma-Aldrich) was added to the immunoprecipitated CDK5 and assays were carried out in reaction buffer (30 mM Hepes, pH 7.2, 10 mM MgCl_2_, and 1 mM DTT) containing [γ-^32^P] ATP (10 Ci) at room temperature for 1 h and then terminated by adding SDS sample buffer and boiling for 5 min. Samples were subjected to 15% SDS-PAGE, stained by Coomassie Brilliant Blue, and dried, and then phosphorylated histone H1 was detected by autoradiography. The other half of the IP was used for Western blotting to determine the total amount of CDK5 immunoprecipitated from the SCN samples. To quantify the kinase activity at each time point, the following formula was used: ([^32^P] H1/total H1 for each reaction)/CDK5 IP protein.

### Filter-aided *in vitro* kinase assay, phosphopeptide enrichment and mass spectrometry analyses

Filter-aided *in vitro* kinase assays and mass spectrometry analyses were performed essentially as described (Hatakeyama et al., 2019). Briefly, recombinant Cdk5/p35 (Millipore) was incubated with the GST-fused PER2 protein fragment. On 10 kDa MW-cutoff filters (PALL) samples were incubated in kinase buffer containing 50 mM Hepes, pH 7.4, 150 mM NaCl, 0.625 mM DTT, Phostop tablets (Roche), 6.25 mM MgCl_2_, and 1.8 mM ATP at 30°C for 1 h. Samples without ATP were used as negative control. Assays were quenched by 8 M urea and 1 mM DTT. Protein digestion for MS analysis was performed overnight (Wisniewski, Zougman, Nagaraj, & Mann, 2009). Phosphopeptides were enriched by metal oxide affinity enrichment using titanium dioxide (GL Sciences Inc., Tokyo, Japan) (Zarei, Sprenger, Rackiewicz, & Dengjel, 2016).

LC-MS/MS measurements were performed on a QExactive Plus mass spectrometer coupled to an EasyLC 1000 nanoflow-HPLC. Peptides were separated on fused silica HPLC-column tip (I.D. 75 µm, New Objective, self-packed with ReproSil-Pur 120 C18-AQ, 1.9 µm [Dr. Maisch, Ammerbuch, Germany] to a length of 20 cm) using a gradient of A (0.1% formic acid in water) and B (0.1% formic acid in 80% acetonitrile in water): loading of sample with 0% B with a flow rate of 600 nL min-1; separation ramp from 5-30% B within 85 min with a flow rate of 250 nL min-1. NanoESI spray voltage was set to 2.3 kV and ion-transfer tube temperature to 250°C; no sheath and auxiliary gas was used. Mass spectrometers were operated in the data-dependent mode; after each MS scan (mass range m/z = 370 – 1750; resolution: 70,000) a maximum of ten MS/MS scans were performed using a normalized collision energy of 25%, a target value of 1,000 and a resolution of 17,500. The MS raw files were analyzed using MaxQuant Software version 1.4.1.2 (Cox & Mann, 2008) for peak detection, quantification and peptide identification using a full length Uniprot Mouse database (April, 2016) and common contaminants such as keratins and enzymes used for digestion as reference. Carbamidomethylcysteine was set as fixed modification and protein amino-terminal acetylation, serine-, threonine- and tyrosine-phosphorylation, and oxidation of methionine were set as variable modifications. The MS/MS tolerance was set to 20 ppm and three missed cleavages were allowed using trypsin/P as enzyme specificity. Peptide, site and protein FDR based on a forwards-reverse database were set to 0.01, minimum peptide length was set to 7, and minimum number of peptides for identification of proteins was set to one, which must be unique. The “match-between-run” option was used with a time window of 1 min. The mass spectrometry proteomics data have been deposited to the ProteomeXchange Consortium via the PRIDE partner repository with the dataset identifier PXD012068.

Project Name: Cyclin dependent kinase 5 (CDK5) regulates the circadian clock

Project accession: PXD012068

Reviewer account details:

Username

Password: DEBd5FKk

### Generation of an antibody against phospho-serine 394

We raised in mouse a specific monoclonal antibody recognizing the phosphorylated form of serine 394 of PER2 in collaboration with GenScript Company. The sequence used for the immunogen preparation was: FDY {pSer} PIRFRTRNGEC. 3 Balb/c mice and 3 C57 mice were immunized with conventional strategies and antisera obtained from those animals were used for the first control experiment performed by *in vitro* kinase assay (Fig. S5C). The positive antiserum was used for the cell fusions. Subsequently, a screening with 16 96-well plates (from 2×10E4 clones) was performed by indirect ELISA, primary screening with phospho-peptide, then counter-screening with non-phospho-peptide. The obtained supernatants were tested by *in vitro* kinase assay in order to screen which one was better recognized the phospho-form of PER2 S394 (Fig. S5D). Finally, 5 selected positive primary clones selected were subcloned by limiting dilution and tested as final antibody (Fig. S5E).

### Statistical analysis

Statistical analysis of all experiments was performed using GraphPad Prism6 software. Depending on the type of data, either an unpaired t-test, or one- or two-way ANOVA with Bonferroni or Tukey’s post-hoc test was performed. Values considered significantly different are highlighted. [p<0.05 (*), p<0.01 (**), or p<0.001 (***)].

## Acknowledgments

We thank Stéphanie Aebischer, Antoinette Hayoz, Cressida Harvey, Naila Ben Fredi, Jean-Charles Paterna (Viral Vector Facility, University of Zürich) and the Bioimage platform (University of Fribourg) for technical support. Funding from the Swiss National Science Foundation (31003A_166682, 310030_166474/1 and 316030_177088) is acknowledged. A.B. was supported by a fellowship from the Fondazione Cenci Bolognetti, Instituto Pasteur.

## Supplemental figures

**Figure S1:**
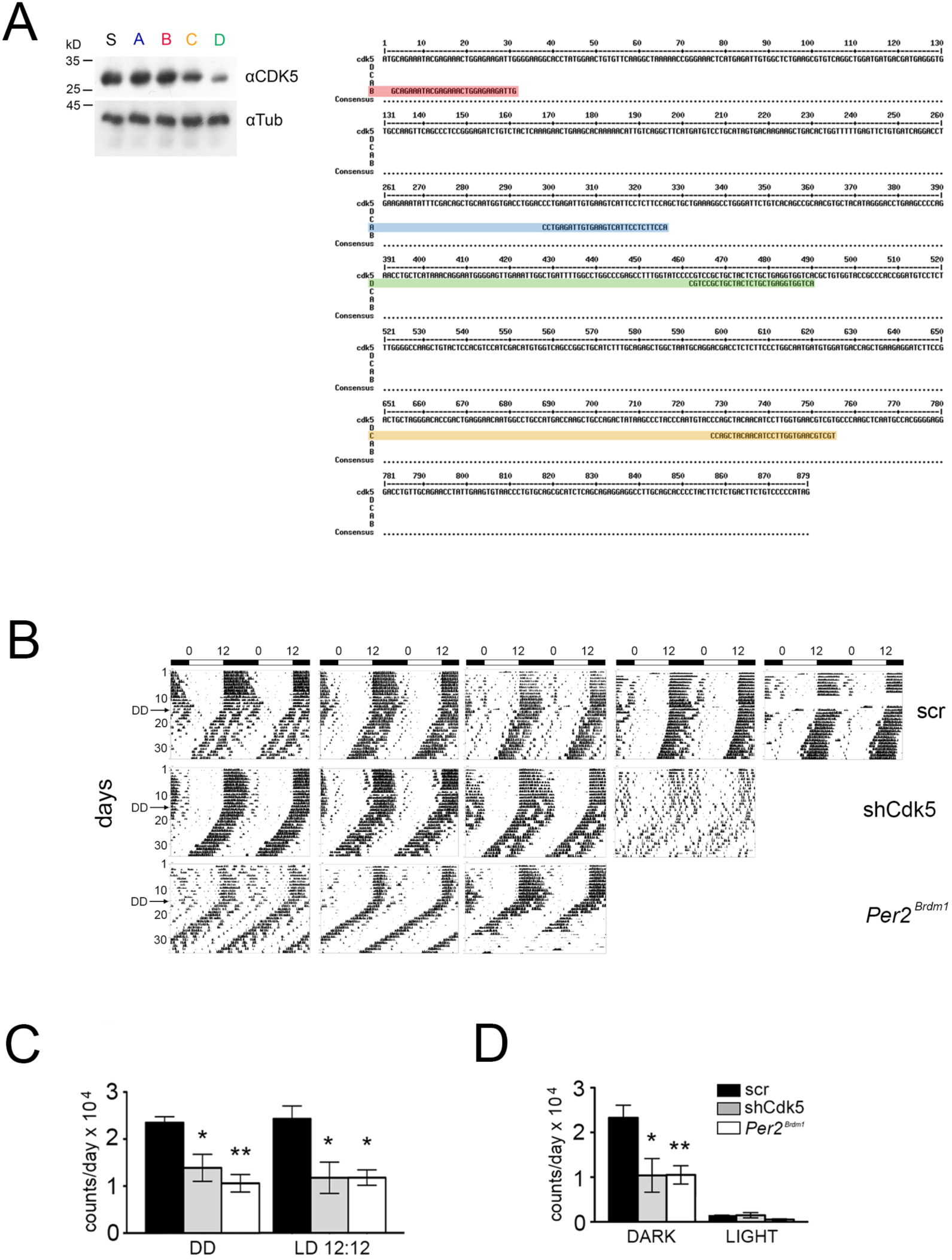
Molecular and behavioral investigation of CDK5 silencing activity in cells and mice. (A) Western blot using NIH 3T3 cell extracts transfected with different shRNAs against *Cdk5*. All shRNAs were mapped to the *Cdk5* sequence. The Western blot reveals that the shRNA D (nucleotides 462 to 490) showed the best silencing activity. (B) Wheel-running activity of mice infected with AAV expressing scrambled shRNA or shCdk5 and animals with a deletion in the *Per2* gene (*Per2^Brdm1^*) used for the statistical analysis in Fig. 1B and Fig. S1 B, C. (C) shCdk5 (13878±2877 counts/day, n=6) and *Per2^Brdm1^* mice (10598±1856 counts/day, n=4) in DD when compared with the control animals (23478±1277 counts/day, n=6) as well as in LD conditions: shCdk5 (11894±3379 counts/day, n=6), *Per2^Brdm1^* mice (11919±1665 counts/day, n=4) and control animals (24577±2787 counts/day, n=6). (D) Dark: scramble (23276±2817 counts/day, n=6), shCdk5 (10399±3764 counts/day, n=6), *Per2^Brdm1^* mice (10521±2052 counts/day, n=4). Light: scramble (1301±223 counts/day, n=6), shCdk5 (1495±582 counts/day, n=6), *Per2^Brdm1^* mice (528±150 counts/day, n=4). (Mean±SEM). 1-way ANOVA with Bonferroni post-test, **p<0.01, *p<0.05.

**Figure S2:**
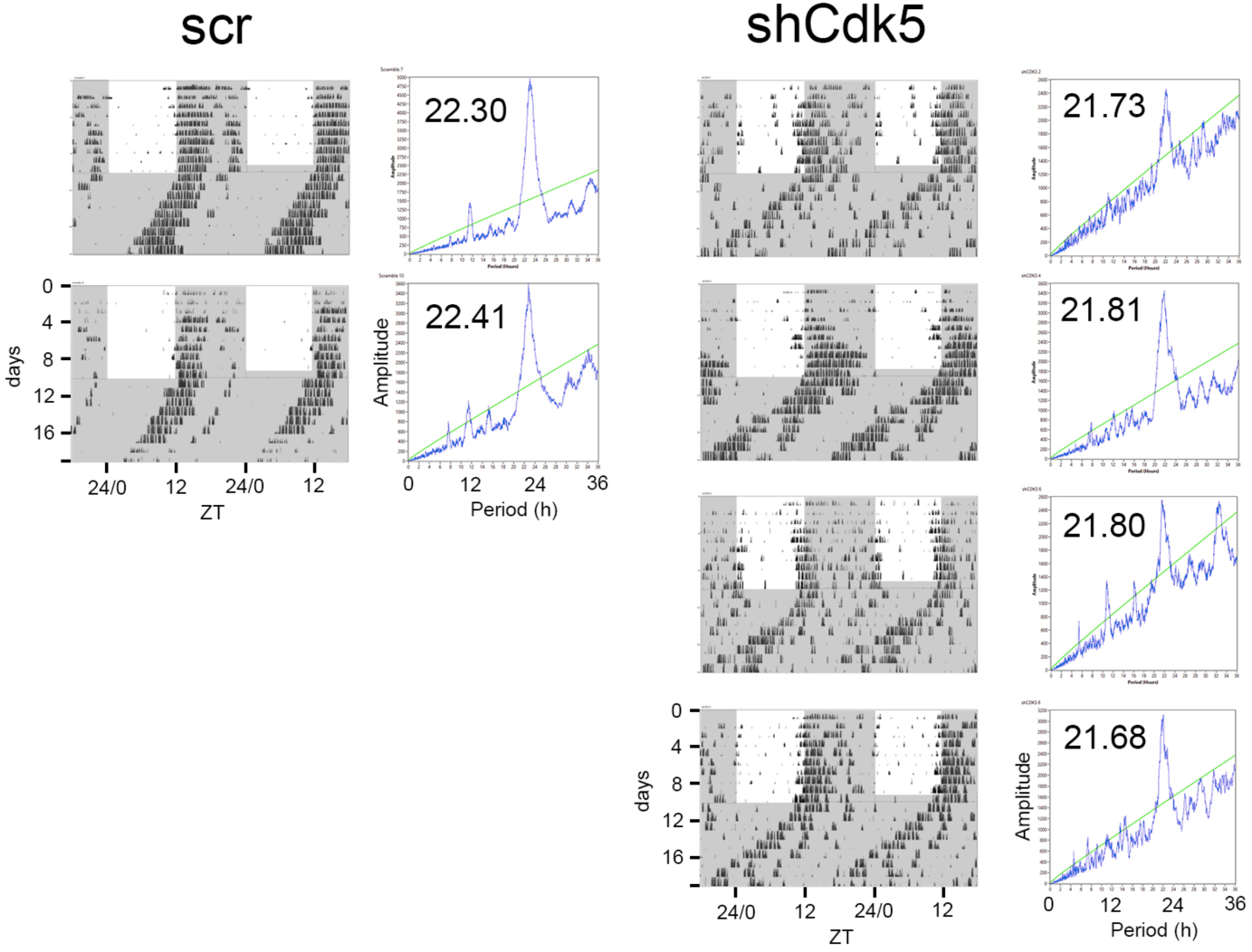
Knockdown of Cdk5 in *Per2^Brdm1^* mutant mice. Wheel-running activity (black bins) of *Per2^Brdm1^* mice infected with AAV expressing scrambled control shRNA (scr), or shRNA against Cdk5 (shCdk5). The actograms are double plotted displaying in one row and below two consecutive days. The dark shaded area indicates darkness during which the free-running period was determined. To the right of each actogram the corresponding χ^2^-periodogram is shown. The number in each periodogram indicates the period of the animal.

**Figure S3:**
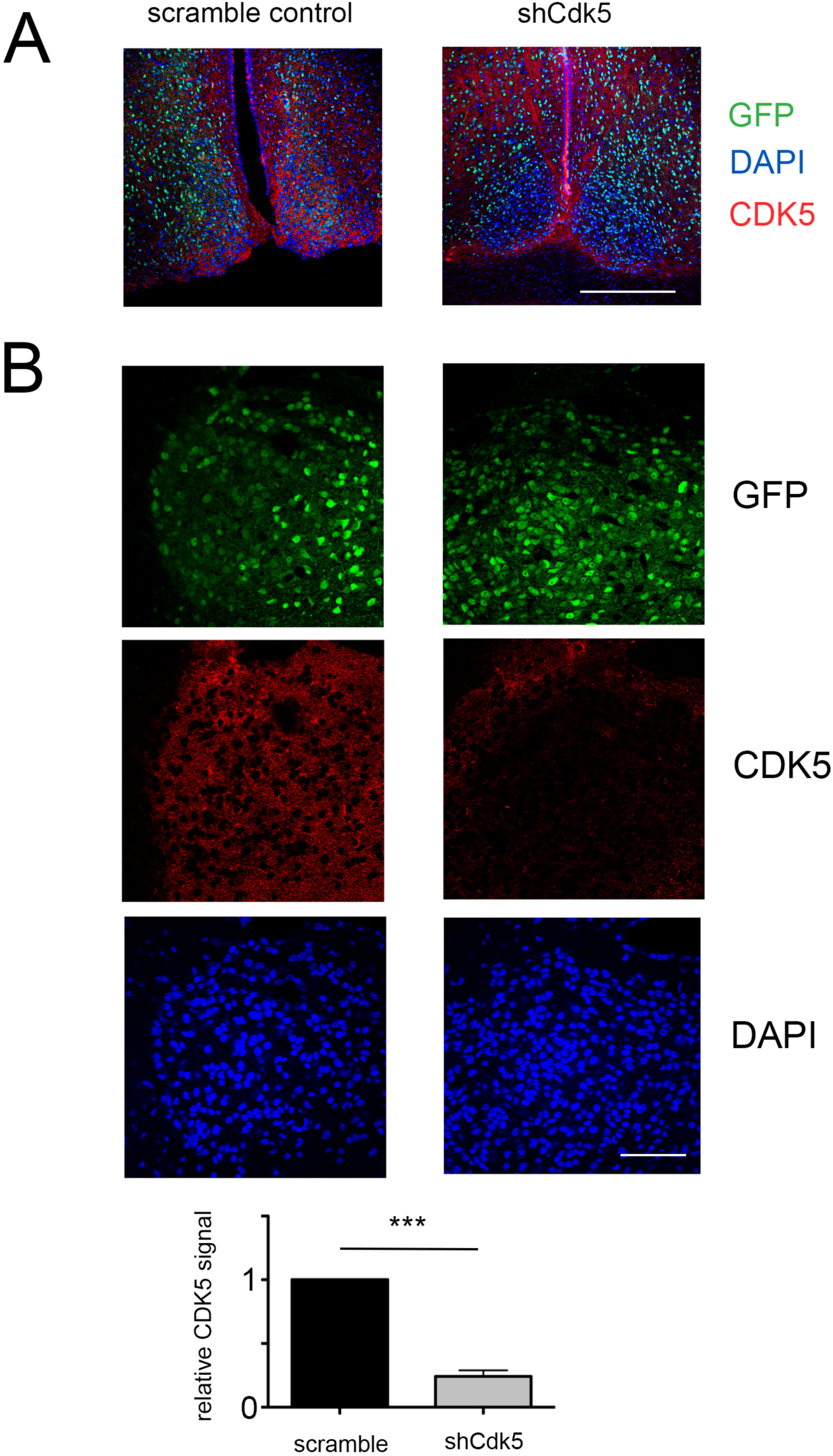
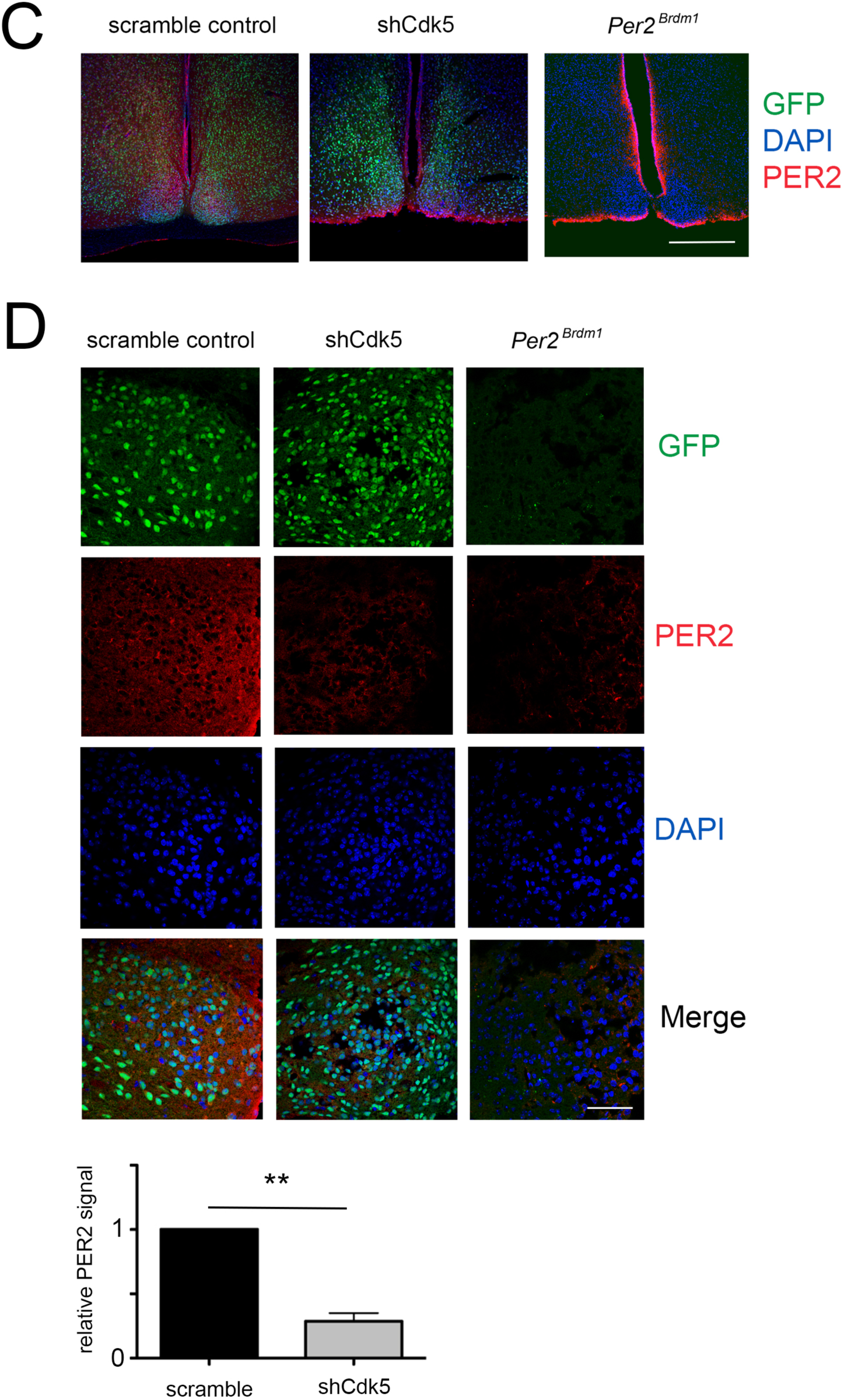
Statistical evaluation of the CDK5 and PER2 signals in the SCN with and without Cdk5 knock-down. (A) Representative brain sections of normal mice containing the SCN region after injection of AAVs carrying either scrambled shRNA or shCdk5. GFP was used as a marker to illustrate the infected region including the SCN. The CDK5 signal (red) is down regulated in the SCN region of AAV shCdk5 injected brain. Scale bar 500 µm. (B) Higher magnification of representative sections of the SCN after AAVs carrying either scrambled shRNA (left column) or shCdk5 (right column). CDK5 if significantly down regulated in brain infected with AAVs expressing shCdk5. Values in the bar diagram represent the mean±SEM of CDK5 signal relative to the signal in the scramble control, t-test, n = 3, ***p<0.001. Scale bar: 60 µm (C) Representative brain sections of normal mice containing the SCN region after injection of AAVs carrying either scrambled shRNA or shCdk5. GFP was used as a marker to illustrate the infected region including the SCN. As control a SCN section of *Per2^Brdm1^* mouse is shown that was not infected with AAV. The PER2 signal (red) is down regulated in the SCN region of AAV shCdk5 injected brain as it was absent in the *Per2^Brdm1^* SCN. Scale bar 500 µm. (D) Higher magnification of representative sections of the SCN after AAVs carrying either scrambled shRNA (left column) or shCdk5 (middle column). CDK5 is significantly down regulated in brain infected with AAVs expressing shCdk5. The right column shows a section of *Per2^Brdm1^* mouse not infected with AAVs. Values in the bar diagram represent the mean±SEM of PER2 signal relative to the signal in the scramble control, t-test, n = 3, **p<0.01. Scale bar: 60 µm.

**Figure S4.**
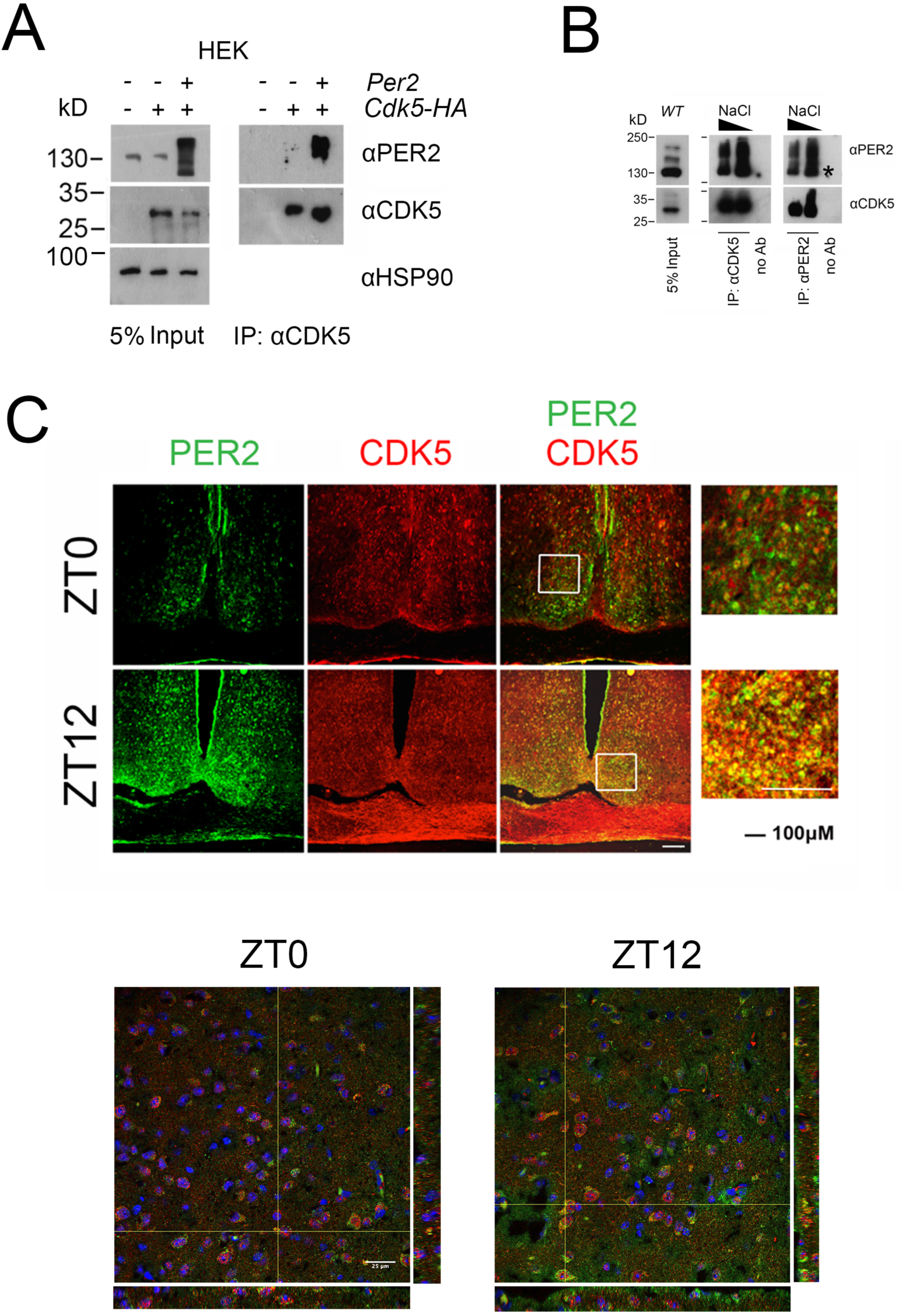
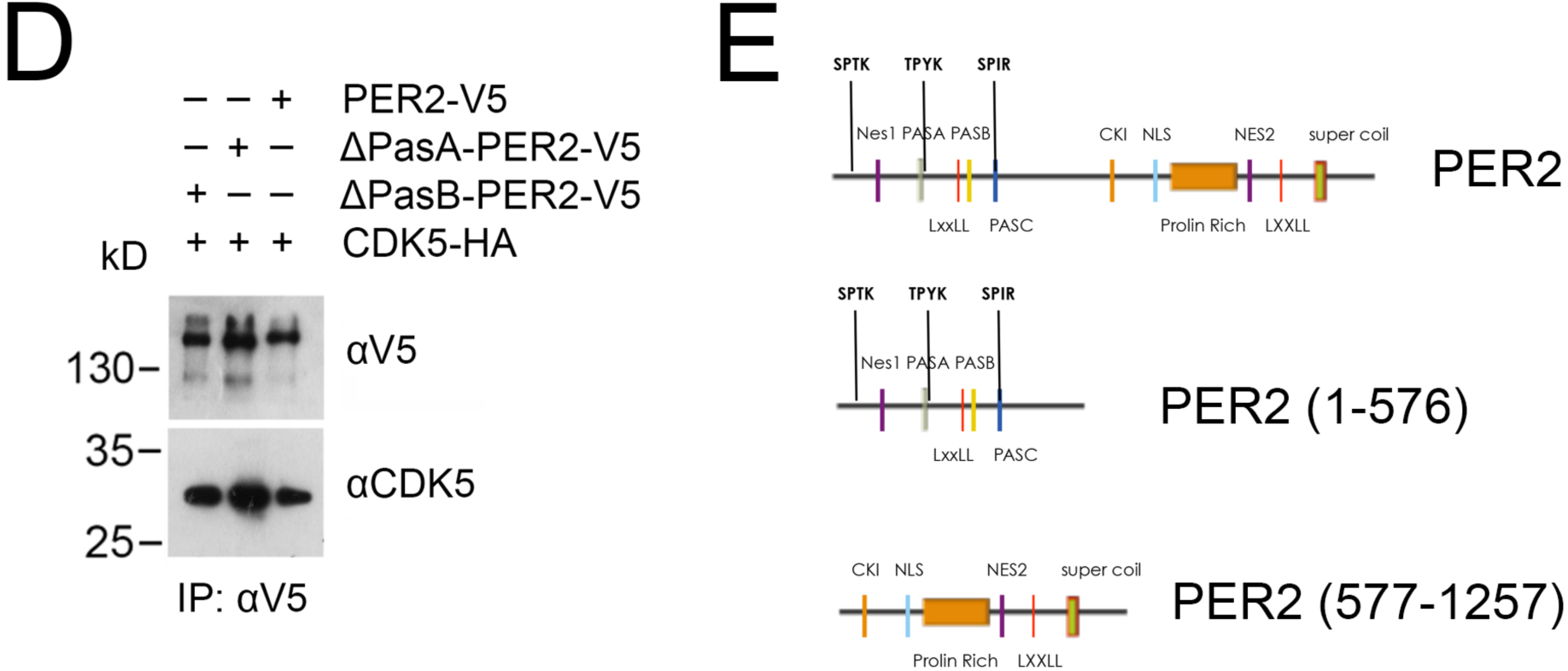
Dynamics of the interaction between CDK5 and PER2. (A) Overexpression of PER2 and CDK5 in HEK 293 cells and subsequent immunoprecipitation (IP) using an anti-CDK5 antibody. The left panel shows the input and the right panel co-precipitation of PER2 with immunoprecipitated CDK5 when both were overexpressed. (B) Immunoprecipitation (IP) of PER2 and CDK5 from total mouse brain extract collected at ZT12. Left panel shows the input. The middle and right panels depict co-immunoprecipitation of PER2 and CDK5 at two different NaCl concentrations using either anti-CDK5 antibody or anti-PER2 antibody for precipitation. * in the blot indicates unspecific signal. (C) Temporal profile of the PER2-CDK5 interaction observed by immunofluorescence at ZT0 and ZT12. SCN slices, obtained from mice perfused at ZT 0 and ZT 12, were stained with anti-PER2 antibody (green) and anti-CDK5 antibody (red). Co-localization of the two proteins results in a yellow color, which was observed only at ZT12. Scale bar: 200 µm. The two panels below show a higher magnification depicting single cells in the SCN. The Z-stacks right and below each image show that PER2 and CDK5 mainly co-localize at ZT12. Scale bar: 25 µm. (D) NIH 3T3 cells were transfected with vectors carrying PER2-V5, ΔPasA-PER2-V5, or ΔPasB-PER2-V5, and subsequently, immunoprecipitation (IP) using an anti-CDK5 antibody was performed. The results showed that CDK5 was able to interact with all forms of PER2. None of the PAS domains of PER2 seems to be involved in the interaction with CDK5. (E) Scheme of PER2 fragments used for the pull-down assy.

**Figure S5.**
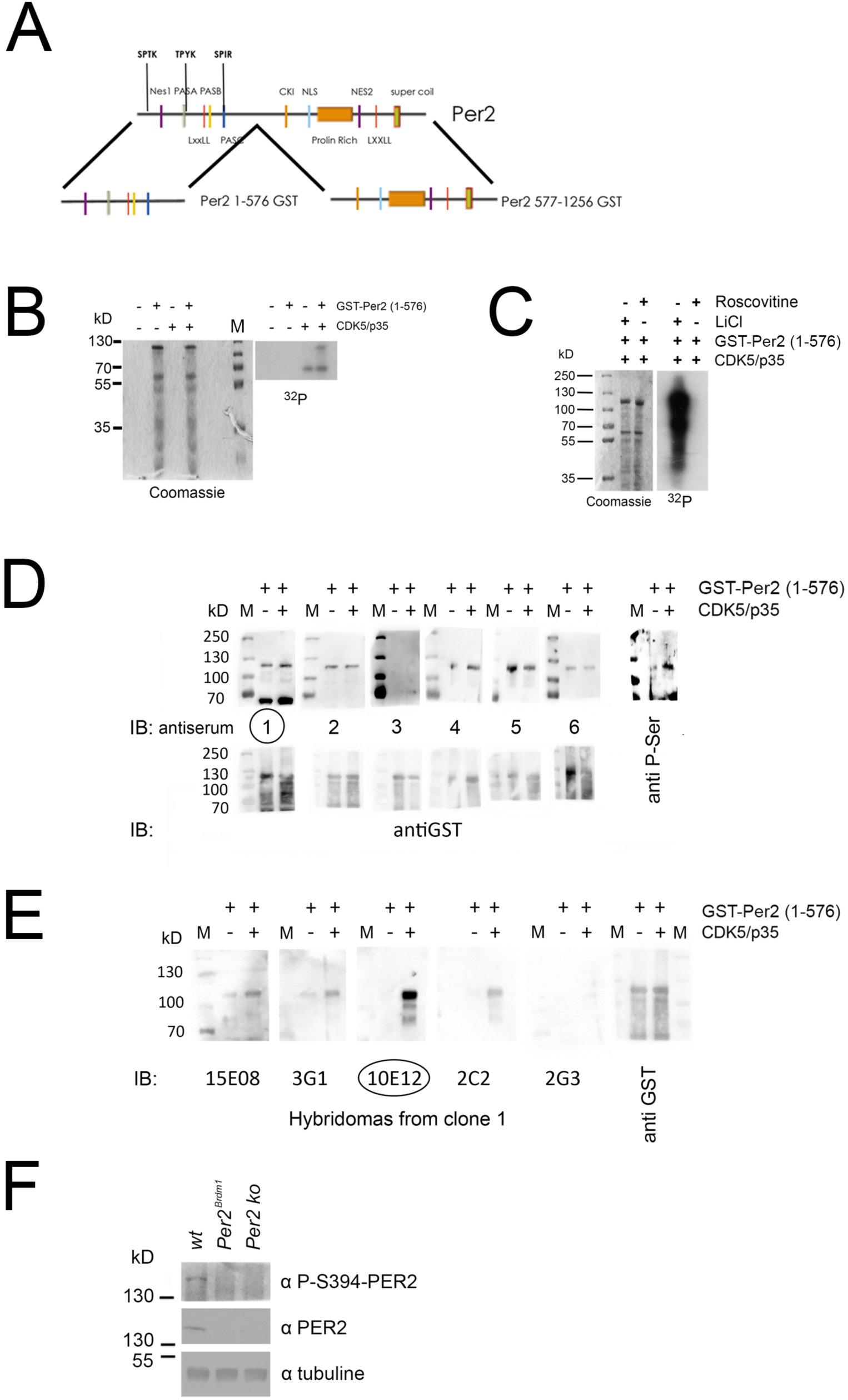
Production and validation of the antibody against the phosphorylated serine 394 on PER2. (A) Scheme of PER2 fragments used for the *in vitro* kinase assay. The fragment 1-576 covers the sites that might be phosphorylated by CDK5 on the basis of the conserved consensus (S/T)PX(K/H/R). (B) An *in vitro* kinase assay performed in presence of recombinant CDK5/p35 and using as substrate the GST-PER2 1-576. (C) The reactions were treated either with LiCl (inhibitor of GSK3β kinase activity) or roscovitine (inhibitor of CDK5 kinase activity) in order to highlight the specificity of the PER2 phosphorylation mediated by CDK5. (D) Different antisera against the PER2 peptide sequence FDY{pSer}PIRFRTRNGEC were tested by *in vitro kinase* assay using recombinant GST-PER2 1-576 (in presence or absence of CDK5/p35) followed by WB. Even if at this stage it was necessary to choose an antiserum that recognized the PER2 peptide regardless of its phosphorylation status, the antiserum 1 was able to discriminate the two forms and was therefore used for the following amplifications. (E) Different hybridomas producing antibodies against the PER2 peptide sequence FDY{pSer}PIRFRTRNGEC were tested by *in vitro kinase* assay using recombinant GST PER2 1-576 (in presence or absence of CDK5/p35) followed by WB. From 16 different clones, the positive ones are shown. The clone 10E12 was used to produce the final antibody. (F) Total protein extracts were obtained from wild-type, Per2*^Brdm1^* and *Per2*^−/−^ mouse brains at ZT12. Western blot was performed in order to validate the specificity of the antibody against the phosphorylated serine 394 on PER2. Only samples obtained from WT tissues showed the phosphorylated form of the protein. Antibody against total PER2 was used as control, which, positively detected the protein only in extracts obtained from wild-type mice.

**Figure S6.**
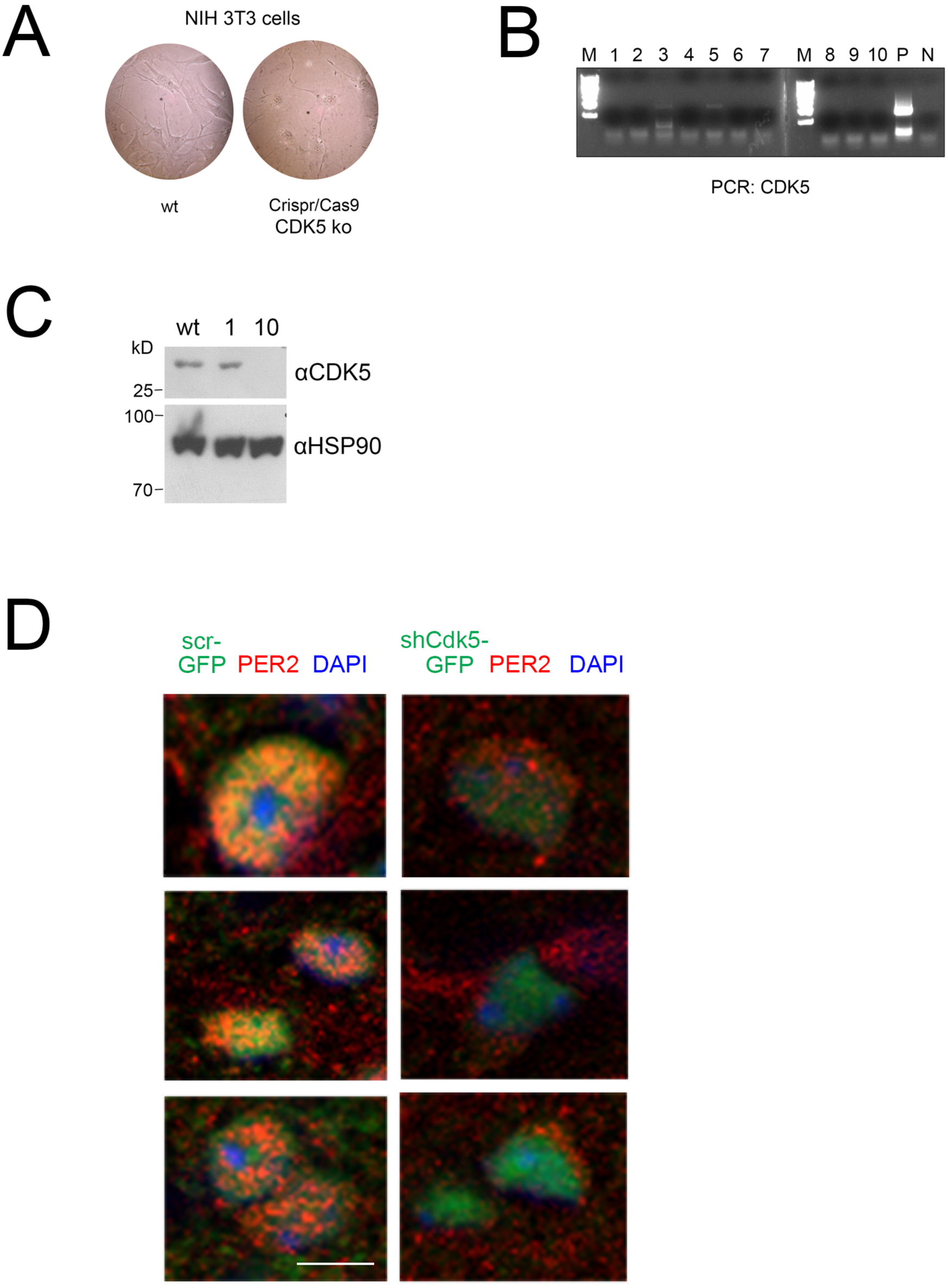
Production and validation of CRISPR/Cas9 *Cdk5*-deficient cell lines. (A) NIH 3T3 and CRISPR/Cas9 *Cdk5*-deficient cells were photographed using a bright light microscope (100 x). A clear difference in shape and thickness between the two cell lines could be observed. CRISPR/Cas9 *Cdk5* cells appeared rather stressed and not to be dividing well. (B) PCR to detect the mutation of the genomic *Cdk5* DNA sequence was performed on different putative knock-out clones. Among these, clones 3 and 5 showed the *Cdk5* PCR product, demonstrating that showed they were false positive for knocking out the gene. A positive control (WT genomic DNA) and negative control (water as template) were used. (C) Total protein extracts were obtained from clone 1, 10 and WT NIH 3T3. Western blot was performed in order to verify which clone no longer expressed CDK5. Clone number 10 was confirmed to be a positive CRISPR/Cas9 Cdk5 knock-out clone. (D) Additional examples of immunofluorescence of PER2 (red) at ZT12 in mouse SCN sections after infection with AAV (green) expressing scrambled shRNA (left column of panels), or shCdk5 (right column of panels). Nuclei are visualized by DAPI staining (blue). PER2 can only be observed in the nucleus in presence but not in absence of CDK5. Scale bar = 2 µm.

**Figure S7:**
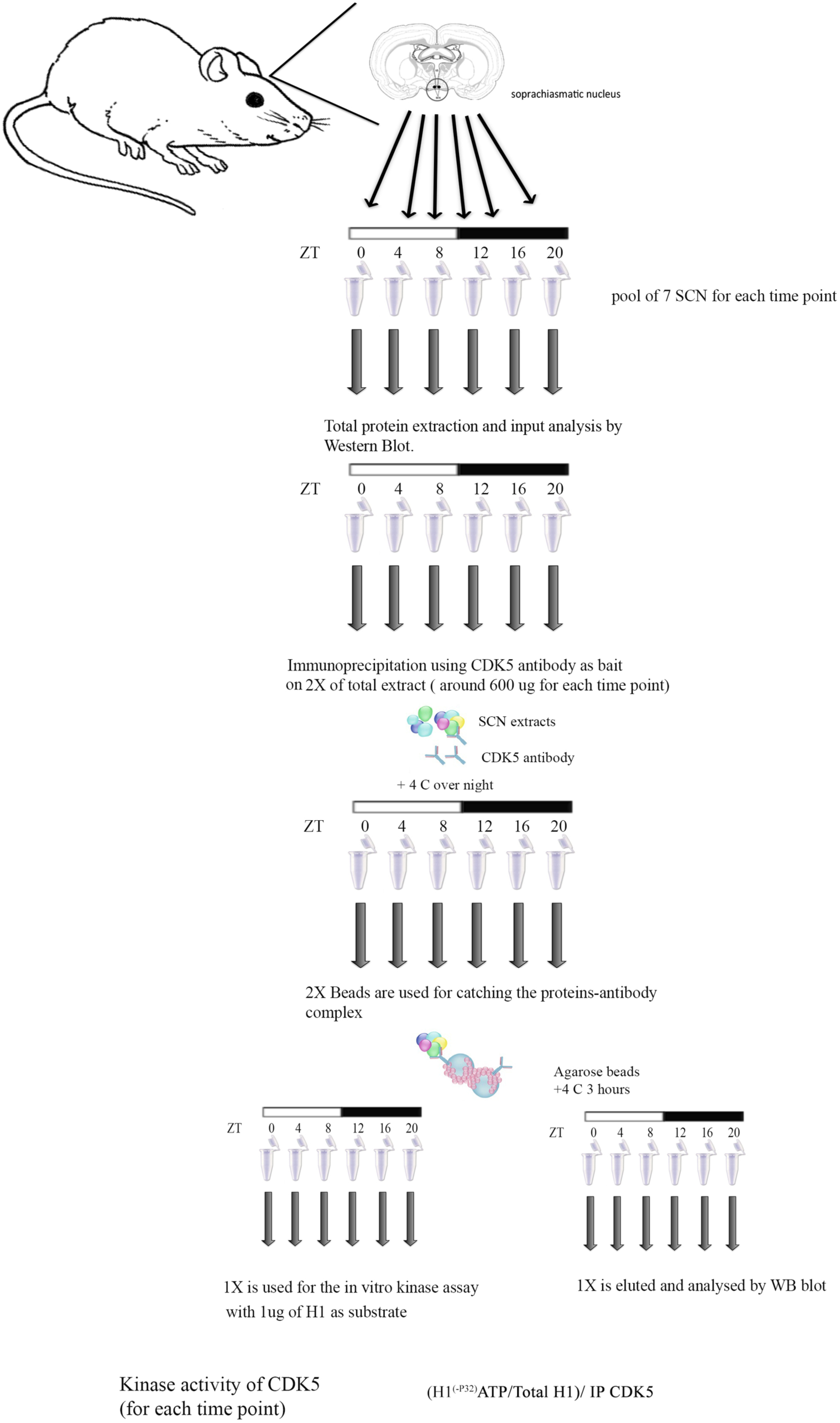
Diurnal CDK5-dependent kinase activity in the SCN. Workflow of the *in vitro* kinase assay performed using immunoprecipitated CDK5 from SCN protein extracts is schematized here. Seven mice were sacrificed, SCN tissues were isolated and pooled together every 4 hours starting from ZT 0 (lights on) until ZT20 (ZT12 lights off). Total protein was obtained from each pool of tissues, the quality of the extracts was checked by WB, and subsequently CDK5 was immunoprecipitated at each time point. Agarose beads detained the immunoprecipitation and one half of the precipitate was used for an *in vitro* kinase assay using as substrate commercial histone H1 as substrate. The other half was analyzed by WB in order to quantify the amount of protein immunoprecipitated, which was used for the kinase assay. Kinase activity around the clock was quantified using the following formula: (^32^P-H1/total H1)/amount of immunoprecipitated CDK5.

## Supplemental Table

**Table S1:**
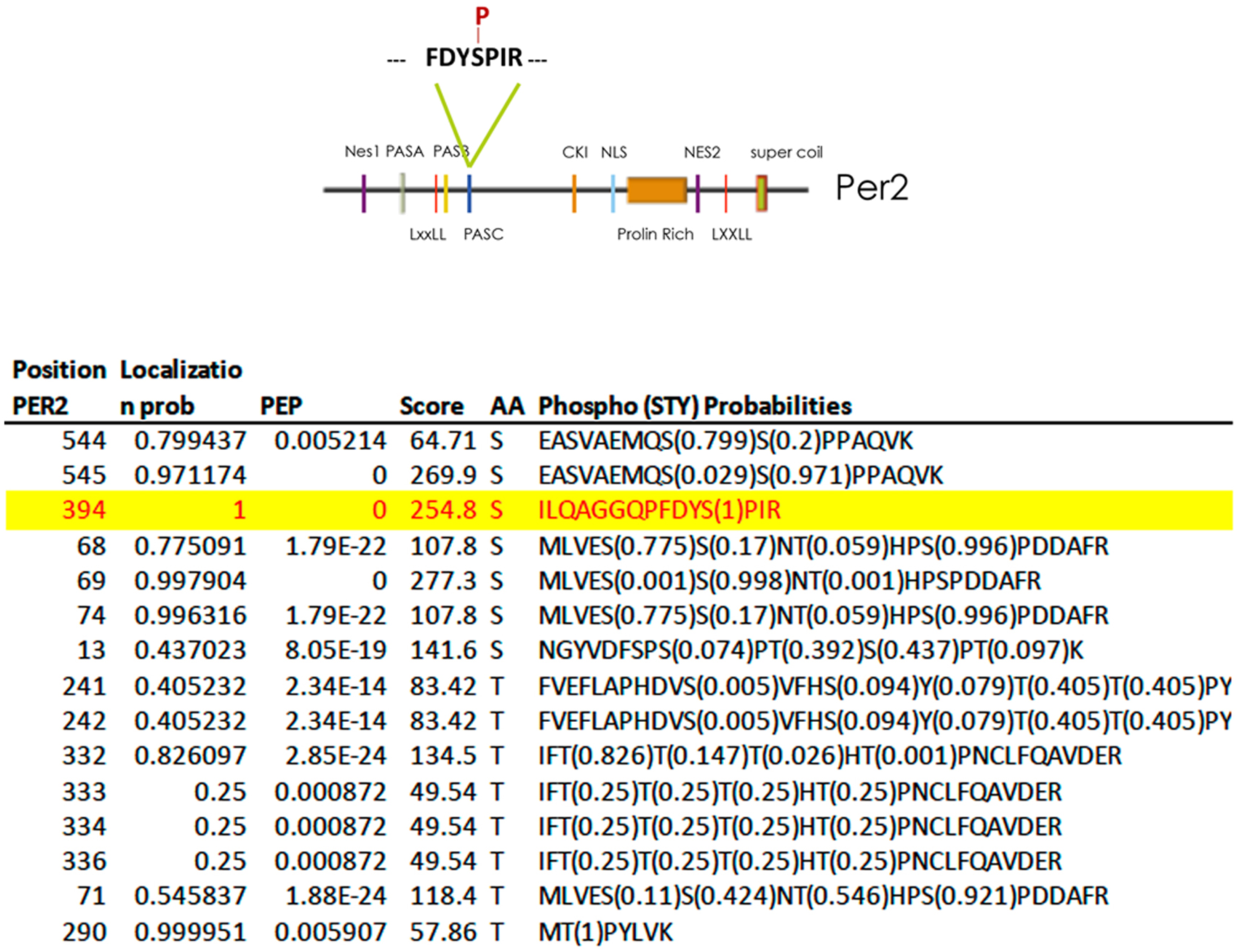
Phosphorylation sites of GST-Per2 (1-576) detected by mass spectrometry. The serine at position 394 stands out as the best localized phosphorylation site within a CDK5 consensus motif with a high peptide score (highlighted in yellow). The colored diagram shows the structural elements of PER2 (1-576) with the S394 phosphorylation site indicated. PEP: posterior error probability; Loc. Prob.; localization probability.

**Table S2:**
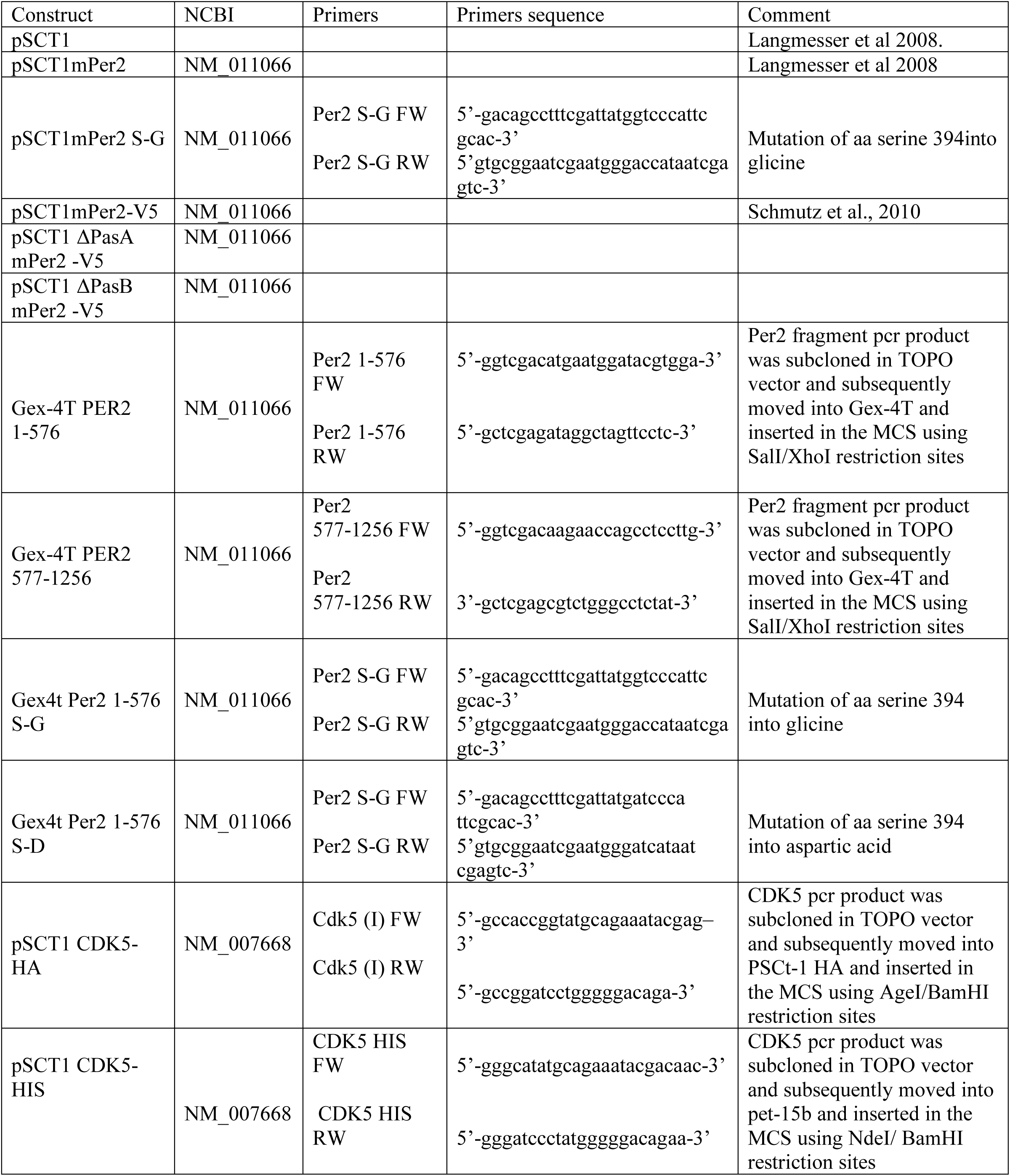
Plasmids.

